# PARP inhibitors enhance reovirus-mediated cell killing through the death-inducing signaling complex (DISC) with an associated NF-κB-regulated immune response

**DOI:** 10.1101/2023.12.21.572541

**Authors:** Joan Kyula-Currie, Victoria Roulstone, James Wright, Francesca Butera, Arnaud Legrand, Richard Elliott, Martin McLaughlin, Galabina Bozhanova, Dragomir Krastev, Stephen Pettitt, Tencho Tenev, Magnus Dillon, Shane Foo, Emmanuel Patin, Victoria Jennings, Charleen Chan, Elizabeth Appleton, Malin Pedersen, Antonio Rullan, Jyoti Choudhary, Chris Bakal, Pascal Meier, Christopher J Lord, Alan Melcher, Kevin Harrington

## Abstract

Oncolytic Reovirus type 3 Dearing (RT3D), is a naturally occurring double-stranded (ds) RNA virus that is under development as an oncolytic immunotherapy We used an unbiased high-throughput cytotoxicity screen of different targeted therapeutic agents with the aim of identifying potential drug-viral sensitizers to enhance RT3D tumour killing. Talazoparib, a clinical poly(ADP)-ribose polymerase 1 (PARP-1) inhibitor, was identified as a top hit and found to cause profound sensitisation to RT3D. This effect was not seen with other classes of oncolytic virus and was not mediated by enhanced viral replication or PARP inhibitor-related effects on the DNA damage response.

RT3D interacts with retinoic acid-induced gene-1 (RIG-I) and activates PARP-1, with consequent PARylation of components of the extrinsic apoptosis pathway. Pharmacological and genetic inhibition of PARP-1 abrogates this PARylation and increases levels of extrinsic apoptosis, NF-kB signalling and pro-inflammatory cell death. Direct interaction between PARP-1 and RIG-I following RT3D/talazoparib treatment is a key factor in activating downstream signaling pathways that lead to IFN-β and TNF-α/TRAIL production which, in turn, amplify the therapeutic effect through positive feedback. Critically, it was possible to phenocopy the effect of RT3D through the use of non-viral ds-RNA therapy and RIG-I agonism. In *in vivo* studies, we demonstrated profound combinatorial efficacy of RT3D and talazoparib in human A375 melanoma in immunodeficient mice. More impressively, in immunocompetent mouse models of 4434 murine melanoma, we achieved 100% tumour control and protection from subsequent tumour rechallenge with the combination regimen. Correlative immunophenotyping confirmed significant innate and adaptive immune activation with the combination of RT3D and PARP inhibition. Taken together, these data provide a clear line of sight to clinical translation of combined regimens of PARP inhibition or ds-RNA agonism, with either viral or non-viral agents, in tumour types beyond the relatively narrow confines of current licensed indications for PARP inhibition.

## Introduction

A number of oncolytic viruses are currently under development as potential anti-cancer therapies^[1-5]^. Drawn from a diverse range of viral species across the spectrum of RNA and DNA genomes, these agents have been extensively tested in wild-type, attenuated and engineered formats in both preclinical and clinical studies. Despite significant effort, the only therapy approved across multiple jurisdictions is a type I herpes simplex virus (HSV), talimogene laherparepvec (T-VEC), for use as a single-agent against melanoma^[3]^. There is a growing consensus that oncolytic viruses will only realise their full therapeutic potential as components of combination treatment regimens. As yet, attempts to augment their efficacy and clinical relevance through combination with standard-of-care therapies, including surgery, chemotherapy, radiotherapy and immune checkpoint blockers have been largely unsuccessful.

In the last two decades, following identification of the BRCA1 and 2 genes and elaboration of their role in homologous recombination (HR)-mediated DNA repair, poly(ADP)-ribose polymerase (PARP) inhibitors have been developed and approved as single-agent, synthetically lethal therapies in patients with HR-deficient breast, ovarian, fallopian tube, peritoneal and prostate cancers^[6-10]^. All existing approvals for PARP1 and 2 inhibitors are predicated on tumours having germline or somatic mutations in components of DNA damage repair pathways and their use is based on specific testing for such abnormalities with companion diagnostic tests^[11]^. Attempts to combine PARP inhibitors with cytotoxic chemotherapy have been significantly complicated by overlapping, on-target toxicities^[12]^.

Previous studies have examined combinations of PARP inhibition with oncolytic HSV^[13]^ or adenovirus^[14]^, with evidence of enhanced anti-tumour effects mediated by viral modulation of DNA repair processes. Such approaches have offered the prospect of inducing a state of PARP inhibitor sensitivity (or BRCAness), but they have not, as yet, translated effectively to clinical trials.

Here, we describe a novel biological interaction between PARP inhibition and the double-stranded (ds) RNA virus, reovirus type 3 Dearing (RT3D, pelareorep), that leads to synergistic anti-tumour activity that is independent of HR deficiency or modulation of DNA repair pathways. Instead, we demonstrate that PARP inhibition affects dsRNA sensing at the level of retinoic acid-inducible gene 1 (RIG-I) with consequent effects on extrinsic apoptosis, NF-kB signalling and pro-inflammatory cell death. Critically, we demonstrate that this effect can be phenocopied by non-viral ds-RNA therapy and RIG-I agonism, providing a clear line of sight to clinical translation of this therapeutic partnership with either viral or non-viral agents in tumour types beyond the relatively narrow confines of current licensed indications for PARP inhibition.

## Results

### Identifying oncolytic reovirus (RT3D) sensitizers

We performed an unbiased, high-throughput small molecule screen in a 384-well plate format in A375 BRAF^V600E^-mutant melanoma cells to discover synergistic interactions between RT3D and 80 different small molecule inhibitors, approved or in late-stage development for cancer treatment (**Supplementary Table 1**). In brief, A375 cells were incubated with small molecule inhibitors in triplicate for two hours prior to infection with RT3D or control. After 72 hours, cell viability was assessed by a luminescent assay. The effects of combining eight different concentrations of each small molecule inhibitor (0.5-1000 nM) with a range of multiplicities of infection (MOIs) of RT3D (0.01-5.0) were assessed by calculating drug effect (DE) robust Z scores, with values below -2 representing a profound sensitisation effect ^[15, 16]^.

### RT3D combined with the clinical PARP inhibitor talazoparib exerts a robust synergistic effect *in vitro* and *in vivo*

Talazoparib, (Lead therapeutic, Pfizer), an approved poly(ADP)-ribose polymerase 1/2 (PARP-1/2) inhibitor^[17]^ caused profound sensitisation to RT3D in the high-throughput screen (**Fig. 1A, Supplementary table 2**). To validate this finding, RT3D plus talazoparib was tested on a panel of melanoma cell lines with different genetic backgrounds (BRAF mutant, RAS mutant and wild-type BRAF/RAS), to determine if virus-drug synergy persisted in multiple molecular contexts. Importantly, combination with talazoparib enhanced the effects of RT3D, regardless of the tumour cell line used, suggesting a robust synthetic lethal cytotoxic effect that remained irrespective of treatment schedule (**Fig. 1B, S1, S2A**). This effect was significant and supra-additive when assessed by Bliss independence analysis (**Fig. S2B**). This effect is depicted for A375 and MeWo cell lines using the SRB assay (**Fig. 1C**) and propidium iodide/Hoechst cell death assay with effects dependent on drug concentration and viral MOI (**Fig. 1D, E**). In addition, the effect of RT3D and talazoparib in triple negative breast cancer (TNBC) cell lines was assessed and a significantly enhanced effect was seen in the SUM149, Cal51(human cell line) as well as in the 4T1 mouse cell line compared to either agent alone (**Fig. S2C**).

**Fig 1:**
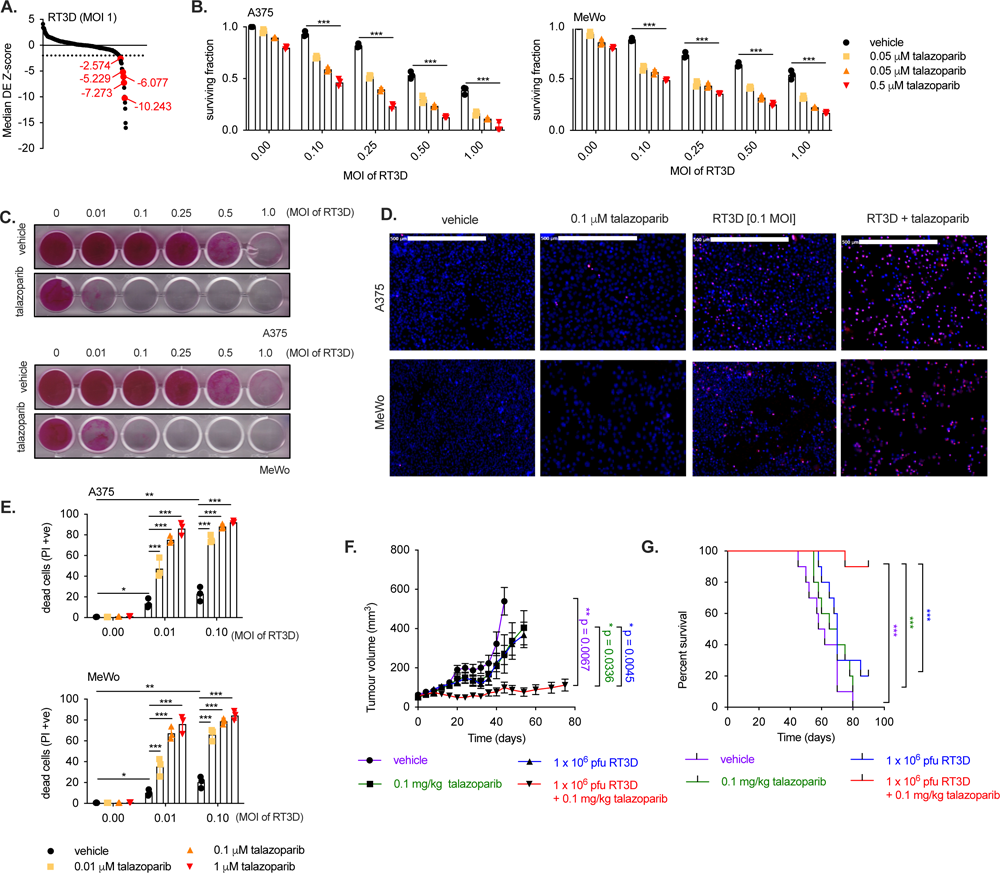
RT3D combination with the PARP inhibitor talazoparib exerts a synergistic effect. **A.** Results from a high-throughput screen experimental set up in the A375 melanoma cell line showing the RT3D multiplicity of infection (MOI) dose of 1 and a range of different chemical inhibitors (ranging from 0.5-1000nM). Cells were analysed 72 hours post-infection using the Cell Titer Glo (CTG) Assay. The z score (DE effect z score = virus effect z score) is plotted on the y axis and was obtained from the standardised value from (i) the median of triplicate samples normalised to virus only versus untreated (ii) the mean of virus only versus untreated and (iii) the median absolute deviation between (i) and (ii) - also known as Z-MAD. **B.** A375 and MeWo cells were pre-treated with increasing doses of talazoparib (0.005 μM, 0.05 μM or 0.5 μM) and thereafter infected with increasing doses of RT3D (MOI of 0,1, 0,25, 0.5 and 1). Cell survival was assessed by MTT at 72 hrs post-infection as represented. **C.** Cell survival was measured using the SRB assays following treatment with 0.1 μM of talazoparib and increasing doses of RT3D at 72 hours post-infection. **D.** A cell death assay was used to measure the uptake of propidium iodide (PI) following RT3D and talazoparib treatment at 48 hours post-infection in A375 and MeWo cells. Representative pictures of cell death assay where dead cells are shown for PI uptake (red) and nuclear staining by Hoechst (blue). **E.** Propidium iodide (PI) uptake following RT3D and BMN 673 treatment at 48 hours shown from 3 independent experiments. (* = p value, **= p value <0.01 and *** = p value <0.001). **F.** CD1 nude mice carrying A375 tumour xenografts were treated with oral administration of vehicle or 0.1 mg/kg talazoparib from Day 1-5. RT3D was injected intratumorally on Day 3 at 1x10^6^ pfu, or sham injection. Sizes of tumours were measured for each treatment cohort, n=10 per group, measured by one-way ANOVA. **G.** Kaplan-Meier curve was evaluated for each treatment group to assess the median survival rate. There was significant prolongation of survival in the combination of RT3D and talazoparib compared to either agent alone (***p = 0.0003 using a Log rank [Mantel.Cox] test).

To assess the wider potential of a RT3D-talazoparib combination in an *in vivo* setting, CD1 nude mice bearing A375 tumours were treated with RT3D, talazoparib or the combination. RT3D plus talazoparib significantly attenuated tumour growth (**Fig. 1F**) and prolonged survival (**Fig. 1G**) compared to either the control, RT3D, or talazoparib alone.

### Talazoparib does not enhance cell kill in combination with other types of oncolytic viruses

To investigate whether the synergistic effect of PARP inhibition using talazoparib is specific to RT3D, or widespread to other types of RNA oncolytic viruses, we tested the combination of talazoparib with CVA-21 (Coxsackie) or MG1 (Maraba) and confirmed that talazoparib only enhanced the effect of RT3D (**Fig. S3** and **Fig 1**). We are currently investigating the effect of talazoparib combination with the HSV-1 DNA oncolytic viruses and the study has shown a significant synergistic effect following the loss of HSV-1 induced PARylation in the presence of talazoparib (data not shown).

### Synergistic activity of RT3D plus talazoparib and its effects on DNA damage repair

Nuclear PARP-1 has a role in DNA damage repair (DDR) and the DDR pathway has been implicated in the effects of combining PARP inhibitors and oncolytic viruses ^[13, 14]^. Therefore, we examined the potential role of DDR pathways in mediating synergy between RT3D and talazoparib. First, cell cycle analysis revealed single-agent talazoparib caused accumulation of cells at G_2_/M, consistent with the DNA damage expected from PARP inhibition. In contrast, RT3D infection increased the fraction of cells in S phase. Together, both agents caused a marked increase in the sub-G1 fraction consistent with the observed synthetic lethal effect (**Fig. S4A**). Talazoparib, but not RT3D, induced markers of DNA damage and repair proteins γH2AX and 53BP1 foci in A375 and MeWo cells. The combination of RT3D and talazoparib did not enhance H2AX and 53BP1 anymore than with talazoparib alone (**Fig. S4B-E**). Finally, an alkaline COMET assay confirmed no increase in DNA damage due to single-stranded breaks following combined RT3D-talazoparib treatment (**Fig. S4F**). Taken together, these data suggest that the addition of RT3D to talazoparib did not overtly increase the level of DNA damage and, thus, an alternative mechanism must explain the observed synthetic lethality between RT3D and PARP inhibition.

### Synthetic lethality is not mediated by enhanced viral replication

We next assessed whether increased viral replication could play a role in the RT3D-talazoparib synergistic effect. We measured viral replication in cells exposed to virus alone or the combination by one-step growth curve and viral plaque assays. Exposure to talazoparib resulted in a non-significant change in viral replication in A375, MeWo and D04 cells (**Fig S5A, B**). Virus protein production was also assessed by immunoblotting for σ3 and μ1C reoviral outer capsid proteins.

Talazoparib exposure increased μ1C production in A375 and MeWo, but not D04 cells. However, σ3 production was unaltered in all cell lines (**Fig. S5C**). Taken together, these data confirm that synergy between talazoparib and RT3D is not explained by increased viral replication.

### Loss of PARP-1 is synthetically lethal with RT3D

Previous studies have found that PARP-1 activation and PARylation of PARP-1-targeted proteins occurs when cells are infected with oncolytic HSV-1^[13]^ and adenovirus (dl922-947)^[14]^. Similarly, we showed that RT3D infection leads to PARP-1 activation and PARylation across A375, MeWo and D04 cells. This effect was inhibited by talazoparib, as shown by both western analysis (**Fig. 2A**) and PAR ELISA (**Fig. 2B**) and was also manifest *in vivo* (**Fig 2C**).

**Fig 2:**
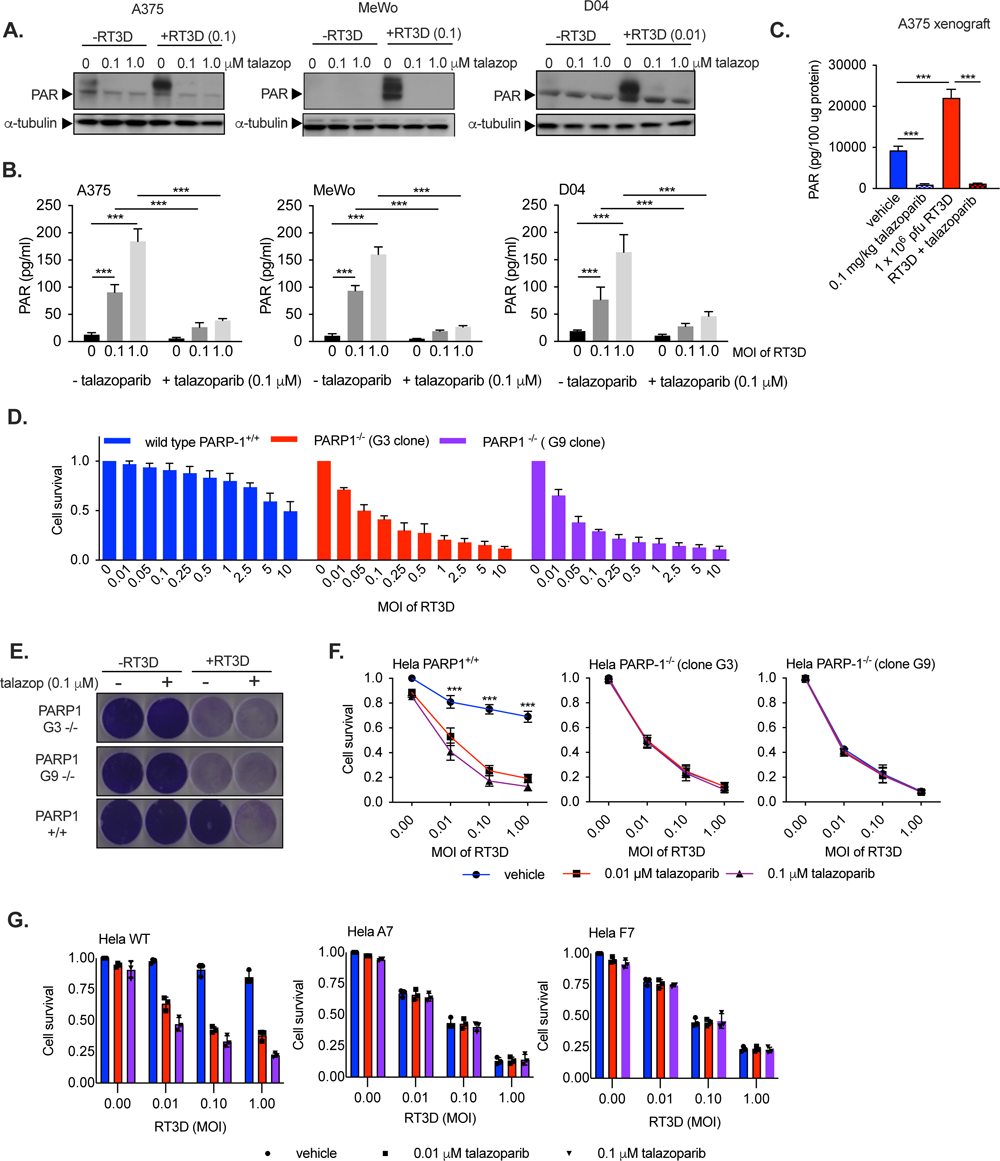
RT3D-induced PARylation is inhibited by talazoparib. **A.** A375, MeWo and D04 melanoma cells were pre-treated with 0.1 μM talazoparib and thereafter infected at an MOI of 0.1 and 1 of RT3D. Cells were harvested, and the lysates collected at 48 hours post-infection. Poly ADP ribosylation (PAR) was assessed by immunoblotting. **B.** PAR was also assessed by ELISA. **C.** CD1 nude mice bearing A375 tumour xenografts were treated with oral administration of vehicle or 0.1 mg/kg talazoparib from Day 1-5. RT3D was injected intratumorally on Day 3 at 1x10^6^ pfu, or sham injection. Following the last treatment on day 5, tumours were harvested and assessed for PAR by ELISA. **D.** RT3D sensitivity was assessed in HeLa PARP-1 paired cells: PARP-1^+/+^ (wild type) and PARP-1^-/-^ (clones G3 and G9), where cytotoxicity was measured by MTT assay 72 hours post-infection. **E.** Cell viability assays were carried out to assess RT3D plus talazoparib in HeLa PARP-1 paired models (PARP-1^+/+^ and PARP-1^-/-^) as shown by crystal violet assays. **F.** SRB cell viability assay to assess RT3D plus talazoparib in HeLa PARP-1 paired models at 72 hours post-infection. Results are shown from 3 independent experiments. * = p value, **= p value <0.01 and *** = p value <0.001. **G.** SRB cell viability assay to assess RT3D plus talazoparib in HeLa PARP-1 mutant models (A7 & F7) versus wild type. Results are shown from 3 independent experiments.

To further elucidate a potentially protective role of PARP-1 against RT3D-induced cell death, we assessed the effect of RT3D on isogenic HeLa cells with or without genetic ablation of the PARP-1 gene (PARP-1^+/+^ wild-type cells or PARP-1^-/-^ clones G3 and G9). PARP-1^-/-^ clones were significantly more sensitive to RT3D (IC_50_ MOI 0.1 and 0.05, respectively) compared to their PARP-1^+/+^ counterpart (IC_50_ MOI >10) (**Fig. 2D**). Genetic loss of PARP-1 in G3 and G9 cells was phenocopied by talazoparib in PARP-1^+/+^ cells following RT3D infection in terms of cell death, as shown by crystal violet and SRB assay (**Fig 2E, F**). Similar effects were also seen in HeLa PARP-mutant cell lines A7 and F7 mutants, which lack efficient PARylation activity, despite expressing PARP1 was previously described ^[18]^ when compared to wild-type (**Fig 2G**).

We also carried out gene silencing of PARP-1, PARP-2 or PARP-3 by siRNA in A375 cells to determine which PARPs were involved in regulating RT3D-induced PARylation. RT3D-induced PARylation persisted despite siRNA targeting PARP-2 and PARP-3, but not PARP-1 (**Fig. S6A**), as expected because PARP1 is the dominant PARP enzyme in cells. Additionally, resistance to RT3D, which could be overcome by talazoparib, persisted following treatment with siRNA against PARP-2 and PARP-3. In contrast, siRNA against PARP-1 completely phenocopied the cell kill observed by the RT3D-talazoparib combination (**Fig. S6B-F**). Taken together these data suggest PARP-1-induced PARylation confers resistance to RT3D, which can be counteracted by talazoparib.

### PARP-1 regulates PARylation induced by RT3D infection and enhances death-inducing signalling complex (DISC)-mediated apoptosis

RT3D has previously been reported to induce apoptosis^[19-21]^. To probe the mechanisms of cell death, a caspase inhibitor screen was used to identify key caspases involved in RT3D plus talazoparib-mediated cell death. Caspase-8 rescued A375 and MeWo cells from RT3D-talazoparib-induced cell death to a greater extent than other caspase inhibitors (**Fig. 3A**). Western blotting confirmed Caspase-8 cleavage in response to RT3D-talazoparib across A375, MeWo and D04 cell lines. Caspase-8 cleavage correlated with caspase-3 and PARP cleavage, and with loss of the anti-apoptotic proteins cIAP2 and c-FLIP_L_ (known inhibitors of caspase-8) (**Fig. 3B**). We next investigated this effect *in vivo*. Western blot analysis of A375 tumours revealed basal and RT3D-induced PARylation was inhibited by talazoparib, as expected. In accord with *in vitro* studies, RT3D-talazoparib combination resulted in increased levels of apoptosis as measured by cleaved PARP, Caspase-8 and -3 in the tumour (**Fig. 3C**). In isogenic (PARP-1^+/+^/PARP-1^-/-^) HeLa cells, an increase in PARylation following RT3D infection in PARP-1^+/+^, but not PARP-1^-/-^ cells, was observed as expected. The RT3D-induced increase in PARylation in PARP-1^+/+^ cells correlated with resistance to apoptotic cell death, as shown by absence of caspase-8 and caspase-3 cleavage. In contrast, cleaved caspase-8 and caspase-3 were observed in PARP-1^-/-^ cells in response to RT3D (**Fig. 3D**). Furthermore, this effect in the PARP-1^-/-^ cells could be phenocopied in PARP-1^+/+^ cells by the addition of talazoparib (**Fig. 3E**).

**Fig 3:**
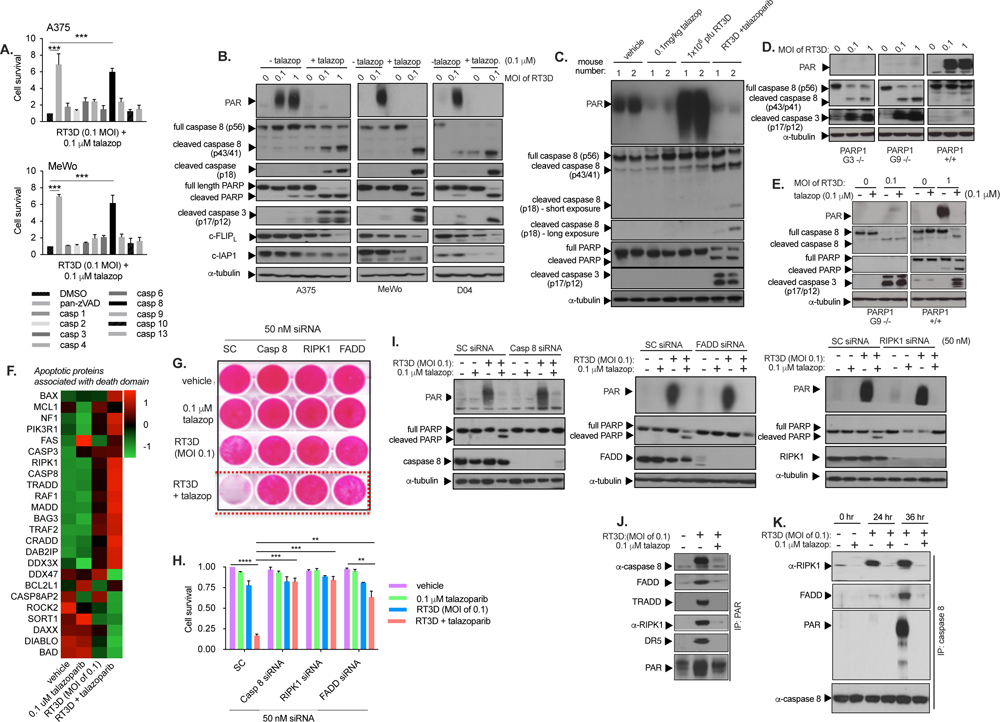
RT3D plus talazoparib enhances death inducing signalling complex (DISC) mediated apoptosis. **A.** A375 or MeWo cells were treated with RT3D (MOI of 0.1) plus 0.1 μM talazoparib in the presence of either pan-Caspase-, or individual Caspase-inhibitors (all at 1 μM) and thereafter measured for cell survival using MTT. **B.** A375, MeWo and D04 cells were treated with 0.1 μM talazoparib and thereafter infected with RT3D (MOI of 0.1) for 48 hours. Western analysis was carried out to assess Caspase-8, Caspase-3 and PARP cleavage and modulation of inhibitors of apoptosis, cIAP1 and c-FLIP_L_. Equal loading of proteins was assessed by probing for α-tubulin. **C.** Western analysis was carried out in A375 xenograft tumours for PAR, Caspase-8, Caspase-3 and PARP cleavage. Equal loading was measured by probing for α-tubulin. **D.** Western analysis was carried out in HeLa PARP-1 paired models. PARP-1^+/+^ (wild type), and PARP-1^-/-^ (clones G3 and G9) were infected with RT3D at MOI of 0.1 and 1.0 and immunoblotted for PAR, Casapse-8 and Caspase-3 cleavage. Equal loading was measured by probing for α-tubulin. **E.** Western analysis was carried out to assess PAR expression and cleavage of Caspase-3 and PARP, following treatment of talazoparib (0.1μM) in HeLa PARP-1^+/ +^ and PARP-1^-/-^ (clone G9) cells. RT3D was then infected in the cells at MOI of• (PARP-1^-/-^ (clone G9) and MOI of 1.0 (PARP-1^+/ +^). Equal loading was measured by probing for α-tubulin. **F.** Proteomic analysis of A375 cells treated with RT3D (MOI of 0.1) in combination with 0.1 μM talazoparib showed upregulation (red) and down-regulation (green) of the apoptotic death domain pathway. Data are an average of 3 independent experiments. **G & H.** A375 cells were transfected with scramble control (SC) or siRNA targeting RIPK-1, Caspase-8 or FADD (all at 50 nM) and subsequently treated with 0.1 μM talazoparib and RT3D (MOI of 0.1) for 48 hours and assessed by SRB cell viability assay. **I.** Western blot analysis was carried out in A375 cells transfected with scramble control (SC) or siRNA targeting Caspase-8, FADD, or RIPKI all at 50 nM and subsequently treated with talazoparib and RT3D at an MOI of 0.1 for 48 hours. Lysates were immunoblotted for PAR, cleaved PARP-1, Caspase-8, FADD or RIPK1 to confirm siRNA target effect. **J.** A375 cells were pre-treated with 0.1 μM talazoparib and infected with RT3D at an MOI of 0.1 and immunoprecipitation (IP) assay with PAR antibody was carried out. Western analysis was carried out to assess the interaction between PARylated proteins and the DISC components (Caspase-8, FADD, TRADD, RIPK1 and DR5). The input (lysate) was carried out to confirm RT3D induced PARylation (refer to supplementary data). Equal loading of proteins was assessed by probing for α-tubulin. **K.** Caspase-8 immunoprecipitation was performed in A375 cells. Z-VAD (10 μM) was added in all samples prior to any treatment to prevent destabilisation of complexes with Caspase-8. Cells were then treated with RT3D (MOI of 0.1)/talazoparib (0.1 μM) at 0, 24 and 48 hours. Western analysis was carried out for PAR, RIPK1 and FADD antibodies. The input (lysate) was carried out to confirm expected RT3D induced PARylation and Caspase-8 cleavage (supplementary fig 7).

To investigate further the mechanistic interaction between RT3D and talazoparib, we analysed the proteome. In support of previous observations implicating caspase-8 in RT3D plus talazoparib-mediated cell death, the proteomic data showed enhanced apoptotic pathways associated with death domain signalling. Apoptotic proteins associated with death domains were assessed. Notably, we saw an increase in the death-inducing signaling complex (DISC), which included Caspase-8 and RIPK1 following RT3D plus talazoparib treatment (**Fig. 3F**).

Subsequently, we silenced components of the DISC, including Caspase-8, RIPK1 and FADD, by siRNA in A375 cells. These conditions protected cells against the combined effect of RT3D plus talazoparib (**Fig 3G, H**). By western blot analysis, RT3D-induced PARylation was not affected by Caspase-8, RIPK1 or FADD siRNA, demonstrating that it occurred upstream of DISC activation (**Fig. 3I**). Conversely, PARP cleavage, as a marker of apoptosis, was prevented by transfection with Caspase-8, RIPK1 and FADD siRNA, implying that the DISC components play an important role in regulating combined RT3D-talazoparib-induced cell death (**Fig. 3I**).

### Talazoparib inhibits RT3D-induced poly(ADP)-ribose interaction with the DISC

Having shown that combined RT3D-talazoparib enhanced apoptosis mediated by the DISC, we explored the possibility of an interaction between DISC components and poly(ADP)-ribose (PAR) chains following RT3D infection. In an immunoprecipitation assay to pull down PAR, we revealed PAR interaction with Caspase-8 (as well as FADD, TRADD and RIPK1). These interactions were abrogated by talazoparib (**Fig. 3J**). Input protein analysis confirmed that RT3D-induced PARylation was inhibited by talazoparib (**Fig. S7A**). Conversely, we immunoprecipitated with pull-down of caspase-8. This revealed an interaction with PAR (by 36 hours) that, again, was prevented by talazoparib (**Fig. 3K**). The interaction of DISC components, RIPK1 and FADD, with caspase-8 was also inhibited by talazoparib and this correlated with an increase in cleaved caspase-8 as shown in the input (**Fig. S7B**). Taken together, these data suggest that RT3D induces PARylation of DISC components to inhibit apoptosis, and that this effect is reversed by talazoparib, leading to enhanced cell death.

### RT3D plus talazoparib-induced cell death is mediated by TRAIL and TNF-∝

TRAIL and TNF-α secretion activate DISC components through TRAIL receptors (DR4) and (DR5) or TNF-α receptors (TNFR1) and (TNFR2), respectively ^[22, 23]^. RT-PCR data showed RT3D-induced increases in both DR4/DR5 (**Fig. S8A**) and TNFR1/TNFR2 expression (**Fig. S8B**) in A375 cells (that were unaltered with talazoparib treatment). RT3D induced TRAIL and TNF-α, measured by ELISA, the effects of which were counteracted by an anti-TRAIL neutralising antibody (2E5) (**Fig. S8C**) or anti-TNF-α neutralising antibody (D1B4) (**Fig. S8D**). Cell death induced by RT3D or combination of RT3D and talazoparib was partially rescued by 2E5-mediated TRAIL (**Fig. S8E**) or TNF-α (**Fig. S8F**) neutralisation. In contrast to blocking TRAIL or TNF-α, activating TRAIL or TNF-α using soluble TRAIL (**Fig. S8G**) or TNF-α (**Fig. S8H**) ligand in combination with talazoparib showed a significant synergistic effect in A375 cells, and this phenocopied the cell death effect seen between RT3D and talazoparib. Like RT3D, TRAIL and TNF-α ligand induced PARylation, and this was lost following talazoparib treatment (**Fig. S8I and Fig. S8J**), respectively.

### NF-κB activity and pro-inflammatory cytokine production is enhanced following RT3D and talazoparib treatment

To evaluate the immunogenic consequences of this combination therapy, a human cytokine array was carried out in A375 cells following RT3D plus talazoparib treatment. The pro-inflammatory cytokines CCL5, CXCL8, CXCL1 and CXCL10 were upregulated (**Fig. 4A**). Next, we evaluated the dependency of these cytokines on DISC components. Silencing of caspase-8, RIPK1 and FADD significantly reduced cytokine production demonstrating the importance of the DISC in regulating cytokine production (**Fig. 4B**). These results correlated with previous data showing that TRAIL-induced cytokine production is regulated through FADD, RPK1 and Caspase 8.

**Fig 4:**
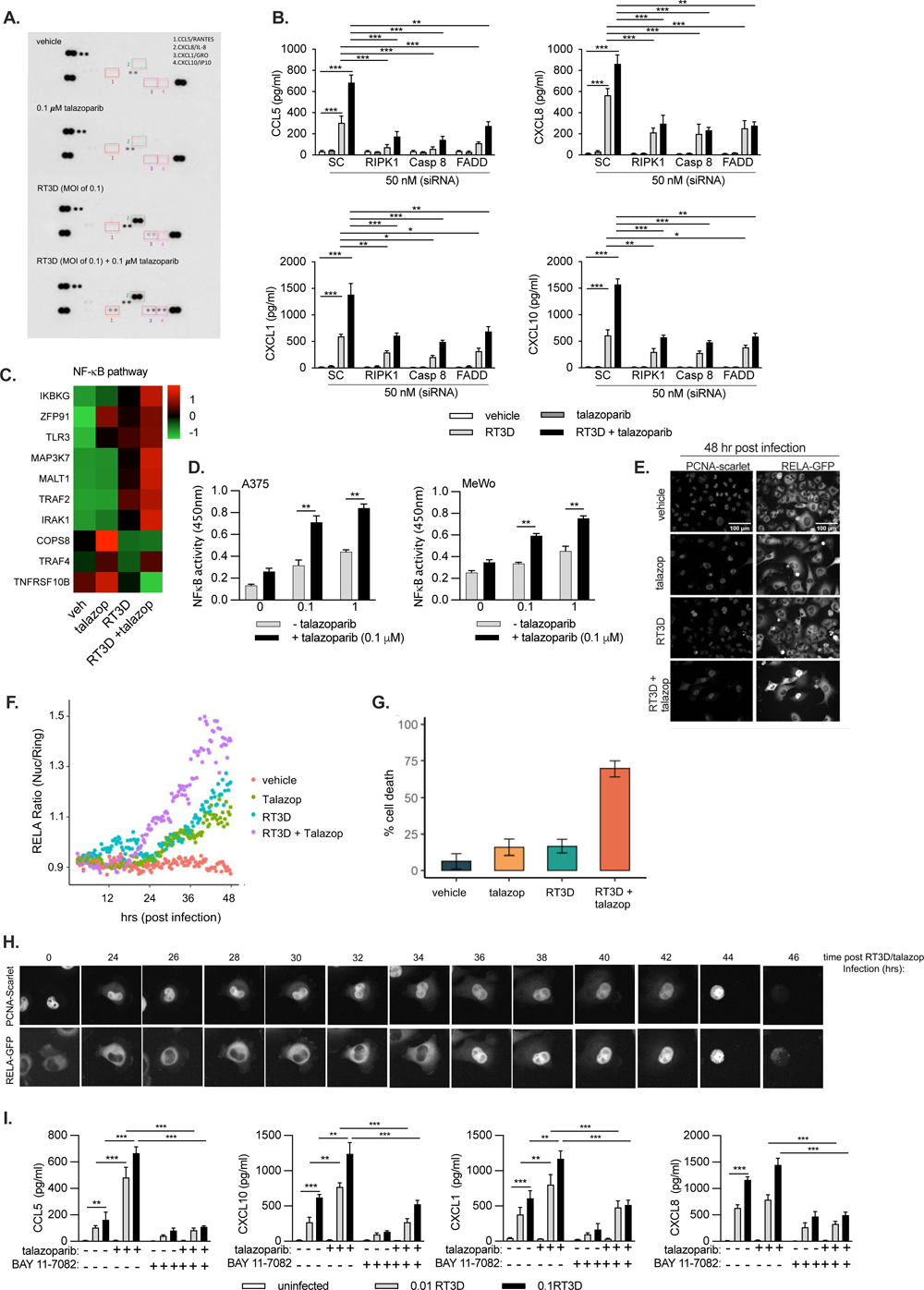
NF-κB activity and pro-inflammatory cytokine production is enhanced following RT3D and talazoparib treatment. **A**. A human cytokine array was used to assess cytokine secretion in A375 following RT3D (MOI of 0.1) and 0.1 μM talazoparib 48 hours post-treatment **B**. A375 cells were transfected with scrambled control (SC), RIPK1, Caspase-8 and FADD siRNA at 50 nM prior to treatment with 0.1 μM talazoparib and RT3D (MOI of 0.1). Supernatants were collected and assessed for CCL5/RANTES, CXCL8/IL8, CXCL1/GRO and CXCL10/IP10 cytokines by ELISA at 48 hours post-infection. All results are shown from 3 independent experiments. **C**. Proteomic analysis of A375 cells treated with RT3D (MOI of 0.1) in combination with 0.1 μM talazoparib to show upregulation (red) and down-regulation (green) of NF-κB-related proteins. Data are an average of 3 independent experiments. **D**. A375 and MeWo cell nuclear extracts were used to assess DNA binding activity of the NF-κB transcription factor RELA (p65) in nuclear extracts following exposure to RT3D (MOI of 0.1) and 0.1 μM talazoparib at 48 hours post-treatment. **E**. Representative images of PCNA-Scarlet (nuclear marker) and Rel-A GFP tagged A375 cells with RT3D (MOI of 10) and 1μM talazoparib treatment. **F**. Rel-A GFP tagged A375 cells were treated with RT3D (MOI of 10) and 1 μM talazoparib over a 48-hour time-period. Cells were imaged by confocal microscopy and single cells tracked using automated imaging analysis. These were then calculated by the nuclear region divided by the ring region RELA-GFP intensity (RELA ratio: summarised as nuclear/ring region intensity) for each timepoint. Data show the mean tracks of RELA ratio over time. **G.** Average percentage of single cell tracks corresponding to dying cells at 48 hrs post treatment. **H**. Representative images of an A375 cell showing high nuclear RELA localisation eventually undergoing cell death following treatment with RT3D (MOI of 10) and 1 μM talazoparib over a 48-hour time-period. RELA-GFP translocates to the nucleus between 34-36 hrs post treatment and cell death is apparent from 44 hrs post treatment. **I.** A375 cells were pre-incubated with the IκBα phosphorylation inhibitor, BAY 11-7082 (5 μM) and then treated with RT3D (MOI of 0.1) plus talazoparib (0.1 μM). Supernatants were collected and assessed for CCL5/RANTES, CXCL8/IL8, CXCL1/GRO and CXCL10/IP10 cytokines by ELISA at 48 hours post-infection.

Proteomics data analysis showed the NF-κB pathway to be upregulated in A375 cells treated with RT3D plus talazoparib, compared to either agent alone (**Fig. 4C**). We performed an ELISA-based assay that specifically detected increased DNA binding activity of the NF-κB p65 transcription factor RELA (p65) in nuclear extracts from cells treated with RT3D plus talazoparib compared to single-agent counterparts, in A375 and MeWo cells (**Fig. 4D**).

To further understand how RT3D plus talazoparib affects NF-κB activity at the single cell level, we tagged RELA with GFP and PCNA with Scarlet at endogenous loci in A375 cells using CRISPR-CAS9 to visualise changes in subcellular localisation of RELA by microscopy, using PCNA as a nuclear marker (**Fig. 4E**). Cells were imaged by confocal microscopy and single cells tracked using automated image analysis. The RELA nuclear:perinuclear cytoplasmic ring region ratio was calculated for each timepoint. The proportion of RELA-responsive cells was highest with RT3D plus talazoparib relative to single-agent counterparts and correlated with the highest percentage of tracked cells dying during the 48 hour imaging period (**Fig. 4F, G**). **Fig. 4H** shows an exemplar cell being tracked following RT3D plus talazoparib treatment where high RELA nuclear localisation was noted around 36 hours prior to cell death, which happened before 48 hours.

We further assessed the synergistic interaction of RT3D and PARP inhibition on cell death and NF-κB signalling using the alternative PARP inhibitors, olaparib and veliparib. RT3D increased NF-κB activity with talazoparib at lower doses when compared with olaparib or veliparib (**Fig. S9A**). Cell viability analysis by crystal violet staining showed that, like talazoparib, olaparib increased RT3D-induced cell death, while veliparib had only modest effects on cell viability (**Fig. S9B**). Taken together, these results confirm that PARP inhibition generally enhances the effects of RT3D on NF-κB signalling, where talazoparib caused a more profound effect compared to other PARP inhibitors. In addition, cytokine induction was increased with olaparib and veliparib, but not to the same extent as with talazoparib (**Fig. S9C**).

To address whether the cytokines upregulated by RT3D plus talazoparib are NF-κB regulated, we used the IκBα inhibitor, BAY11-7082, which is widely used as an inhibitor of downstream NF-κB. RT3D plus talazoparib induced cytokine production of CCL5, CXCL8, CXCL1 and CXCL10 was significantly reduced by BAY 11-7082, highlighting the importance of NF-κB in regulating these cytokines (**Fig. 4I**).

Finally, to assess the link between p65 NF-κB activity and the DISC complex, the DNA binding activity of NF-κB was measured after gene silencing of caspase-8, RIPK1 and FADD. NF-κB activity induced by RT3D or RT3D plus talazoparib was abrogated after siRNA of each DISC component (**Fig. S10A**). Furthermore, NF-κB activity was significantly reduced by co-treatment with neutralising TRAIL antibody (2E5) demonstrating the link between TRAIL and NF-κB signalling (**Fig. S10B**). Collectively, these data suggest that PARP-1 regulates TRAIL-mediated cell death caused by RT3D infection and that this protective effect of PARP-1 can be pharmacologically modulated to elicit enhanced cell death through signalling via DISC and NF-κB pathways.

### Talazoparib enhances the IFN-β signalling pathway through RIG-I

Subsequently, we explored events upstream of NF-κB signalling. Pattern recognition receptors (PRRs) such as the RNA helicase, RIG-I, trigger activation of transcription factors NF-κB and IRF3. Proteomic analysis showed a profound increase in the RIG-I pathway (**Fig. 5A**), and this was further confirmed by western analysis where RIG-I, phosphorylated STAT-1 and phosphorylated IRF-3 were all increased following RT3D plus talazoparib treatment compared to either agent alone (also correlating with a loss of RT3D-induced PARylation) (**Fig. 5B**). The effect of RIG-I signalling on RT3D plus talazoparib-induced cell kill was assessed by silencing RIG-I. This showed that RIG-I siRNA partially rescued enhanced cell kill following RT3D plus talazoparib treatment (**Fig. 5C**). This was confirmed further by western analysis which showed abrogation of PARP cleavage with RIG-I siRNA (**Fig. 5D**). Moreover, RT3D plus talazoparib-induced IFN-β secretion was significantly reduced following RIG-I silencing (**Fig. 5E**).

**Fig 5:**
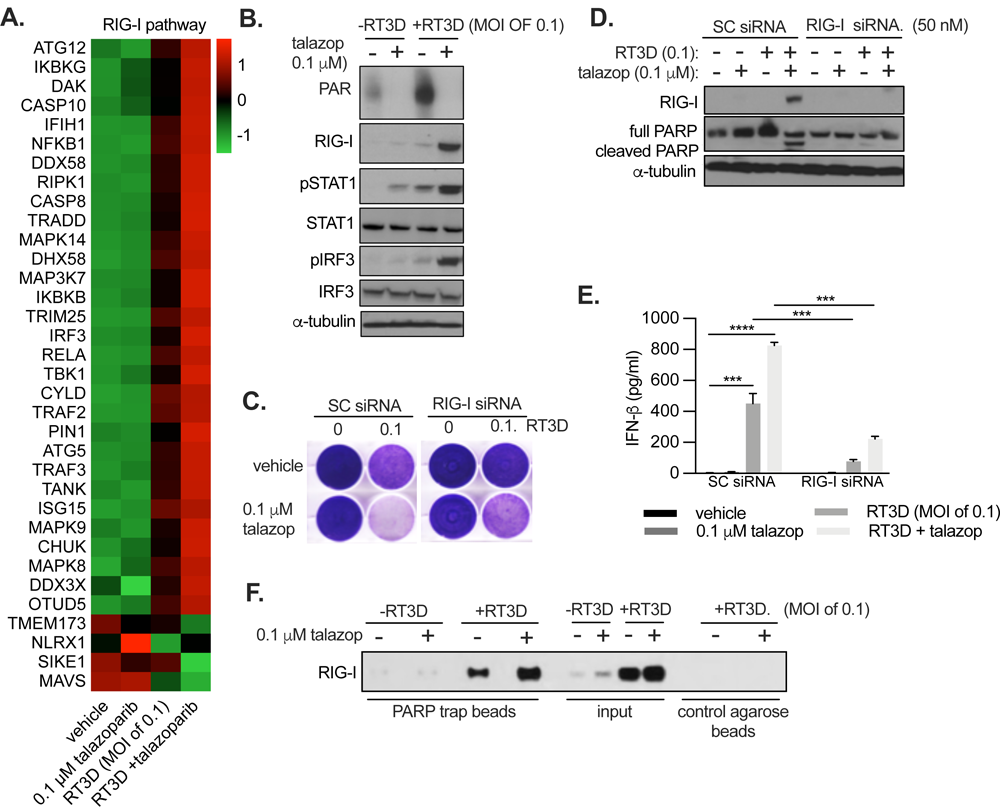
Talazoparib enhances RT3D-induced interferon signalling through RIG-I. **A.** Proteomic analysis of A375 cells treated with RT3D (MOI of 0.1) in combination with 0.1 μM talazoparib to show upregulation (red) and down regulation (green) of RIG-I mediated pathway. Data are an average of 3 independent experiments. **B.** A375 cells were treated with talazoparib and infected with RT3D. Western analysis was carried out to assess RIG-I, pSTAT-1, pIRF3, STAT-1 and IRF3. Equal loading of proteins was assessed by probing for tubulin. **C.** A375 cells were transfected with non-targeting scrambled control (SC) siRNA or siRNA targeting RIG-I (50 nM) and thereafter treated with 0.1 μM and RT3D at an MOI of 0.1 for 48 hours and assessed for cell viability as shown by crystal violet staining. **D.** Western analysis was carried out on cell lysates and probed for RIG-I and cleaved PARP. **E.** IFN-β production was assessed by ELISA. **F.** A375 cells were treated with RT3D (MOI of 0.1) and talazoparib (0.1 μM) and a PARP trap IP was carried out from the lysates. Western analysis was carried out to assess interaction between PARP-1 and RIG-1.

Given the important role of RIG-I in RT3D plus talazoparib-mediated cell death, we tested whether apoptosis induced by RT3D plus talazoparib combination treatment was virus-dependent or whether this effect could also be driven by non-viral RNA sensor agonism. In place of RT3D, we used RNA tool compounds, polyI:C (synthetic dsRNA) and 3p-hRNA (a RIG-I agonist), in combination with talazoparib. Both compounds induced PARylation and phenocopied the effect of RT3D-talazoparib (**Fig. S11A**). Talazoparib inhibited PARylation induced by these tool compounds and that correlated with increased PARP-cleavage (**Fig. S11A**), IFN-β production (**Fig. S11B**) and cell death (**Fig. S11C, D**).

As RNA sensor agonists can trigger PARylation, and since PARP-1 has nucleic acid binding capabilities, we investigated whether PARP-1 interacts directly with RNA sensors like RIG-I using a PARP-1-Trap assay. Our data demonstrated an increase in PARP-1 and RIG-I interaction following RT3D infection and this was increased further by talazoparib treatment (**Fig. 5F**).

### Combination therapy of RT3D and talazoparib enhances anti-tumour efficacy in an immunocompetent animal model

We investigated whether the enhanced effects of RT3D plus talazoparib on apoptosis and cytokine production resulted in increased immunogenicity in an *in vivo* tumour model. Immunocompetent BL/6 mice implanted with 4434 BRAF-mutant melanoma cells were treated with RT3D, talazoparib or the combination (**Fig. 6A**). RT3D plus talazoparib led to a delay in tumour growth (**Fig. 6B**), and prolonged survival (**Fig. 6C**) versus single-agent counterparts. Next, we assessed memory response in the cohort of mice cured following RT3D plus talazoparib treatment by re-challenging with 4434 cells on the contralateral flank. No tumour growth occurred in this cohort of mice (whilst, in parallel, naïve mice developed large tumours over time) (**Fig. 6D**).

**Fig 6:**
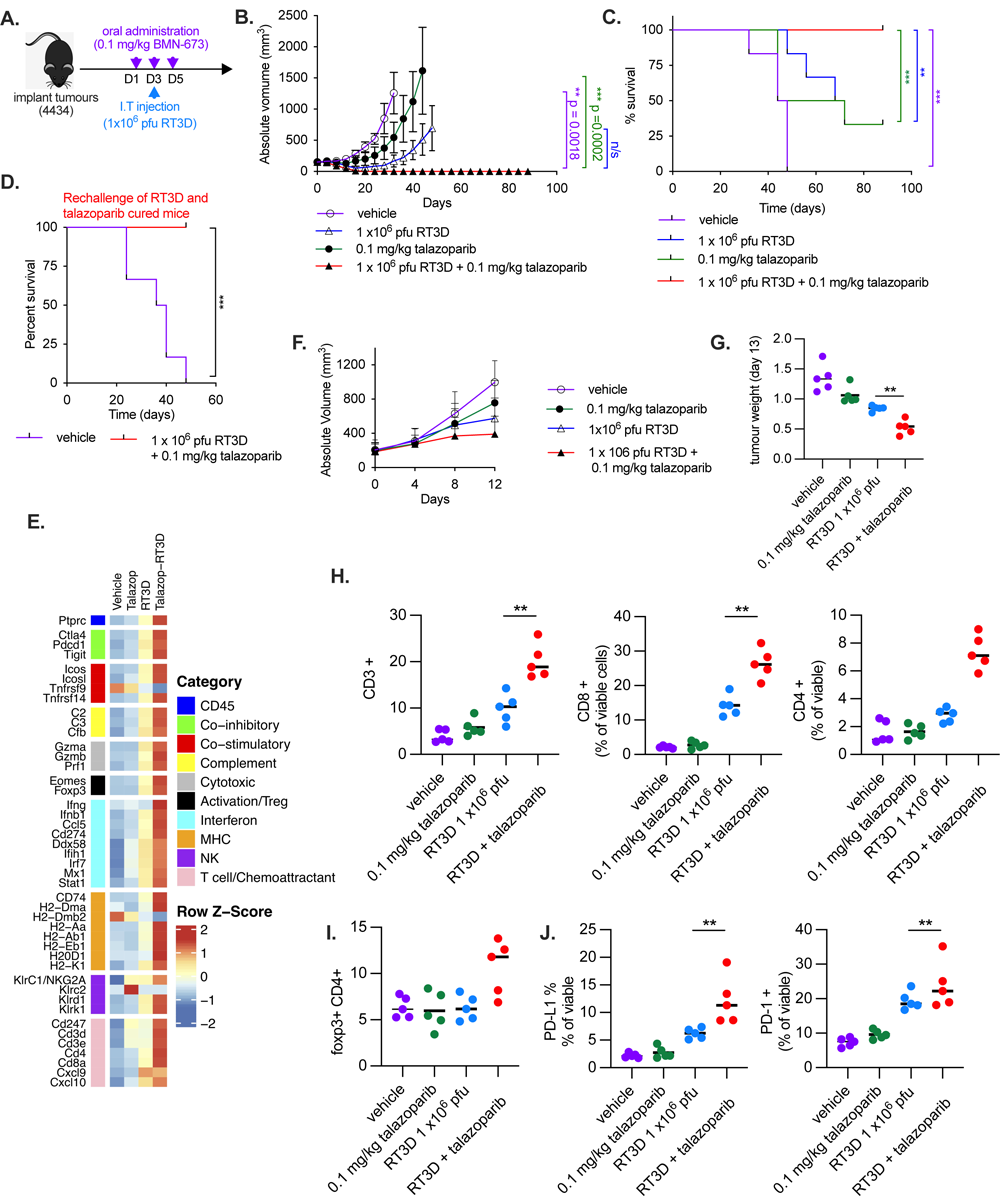
RT3D plus talazoparib enhances anti-tumour efficacy which correlates with an increase in immune response. **A.** Treatment schedule, black/6 mice carrying 4434 tumours were treated with oral administration 0.1 mg/kg talazoparib or vehicle from Day 1-5. RT3D was injected intratumorally on Day 3 at 1x10^6^ pfu or sham injection. **B.** Size of tumours were measured for each treatment cohort consisting of vehicle, talazoparib, RT3D or combination. Each bar represents mean SE± of 10 replicates. **C.** Kaplan-Meier curve was evaluated for each treatment group to assess the median survival rate. **D.** Mice cured following RT3D plus talazoparib (day 90) were rechallenged on the other flank and compared with naïve mice injected with 4434 tumours (both implanted at 4x10^6^ cells) and tumor growth assessed. **E.** Deconvolution of immune cells in 4434 tumours following RT3D and talazoparib treatment. Tumours were dissected on day 8 after treatment, total RNA was isolated and gene expression analysis performed using the mouse Immunology Profiling Panel from NanoString Technologies. **F.** Tumour volumes of mice used for gene expression analysis up to time of harvest (Day 11) were measured. **G.** Tumour weights of mice used for gene expression analysis at time of harvest were measured. **H.** FACS analysis of *in vivo* tumour samples. Data show cell counts of CD3+, CD8+ and CD4+ cells gated from viable cells. **I.** Cell counts of foxp3+ cells gated from viable cells. **J.** Cell counts of PD-L1+ and PD-1+ cells gated from viable cells.

**Fig 7:**
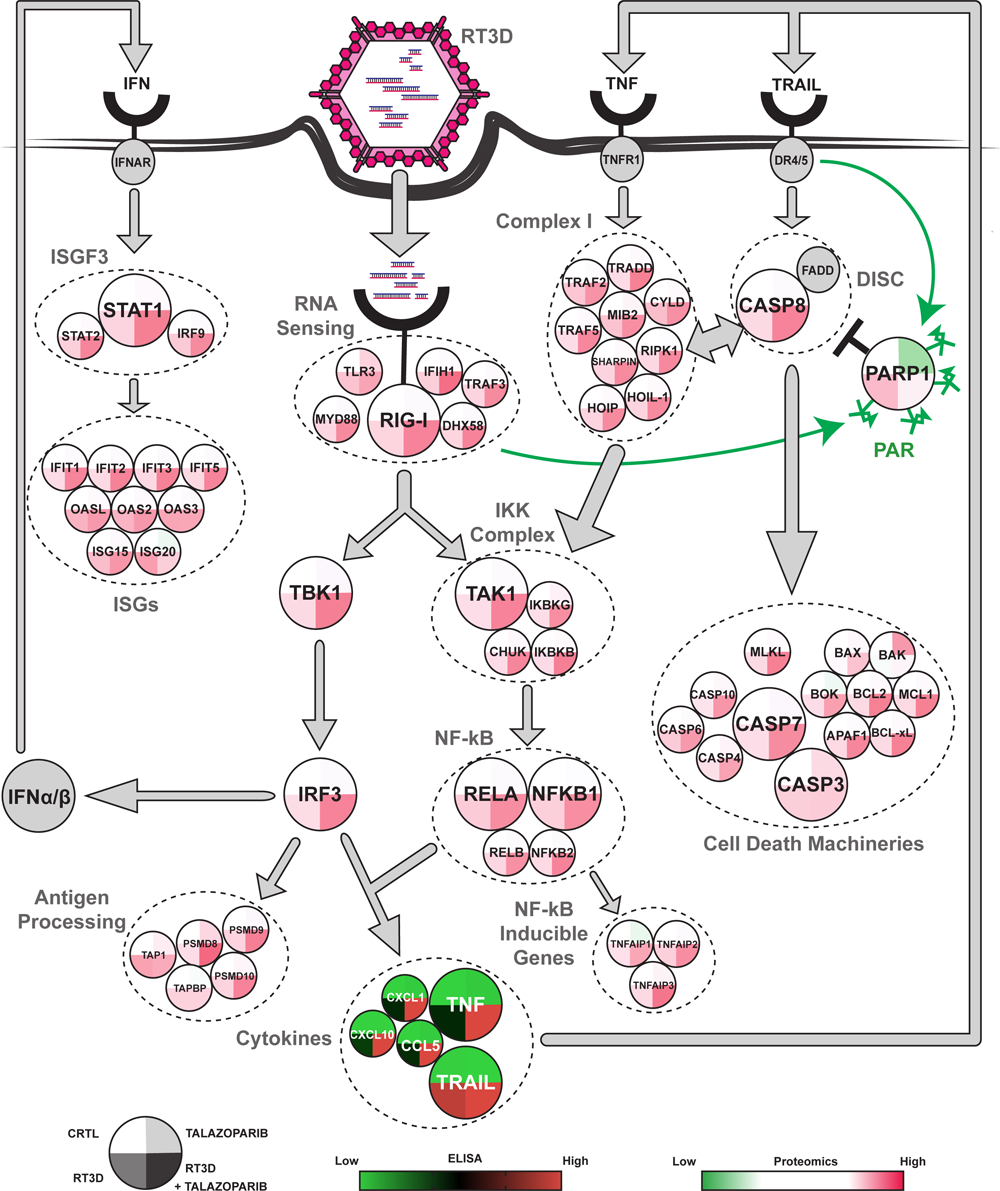
Graphical representation of proteomic and cytokine secretion analysis. Depiction of relevant proteins involved in the recognition, signalling and execution of RT3D infection and talazoparib treatment by mass spectrometry. Each protein is represented by a circle divided in four parts: vehicle, talazoparib, RT3D and RT3D/talazoparib. These levels are color-coded from green (low expression) to white (identical expression) to red (high expression) and represent their value normalised to vehicle. In addition, cytokines that were analysed by ELISA are summarised.

To probe further the impact of RT3D plus talazoparib therapy *in vivo*, total RNA was isolated and gene expression analysis was carried out in the tumour cells using the Nanostring nCounter technology platform where deconvolution was carried out to characterise the immune pathways that were modulated following RT3D and talazoparib treatment. There were increases in tumour-infiltrating immune cells in the microenvironment by means of increased expression of transcripts associated with immune subsets (**Fig. 6E**). In support of these findings, FACS analysis of *in vivo* tumour samples revealed an increase of CD3+, CD8+ and CD4+ cells with RT3D plus talazoparib (**Fig. 6F-H**). Whilst there was an increase in foxp3+ cells with RT3D injection, these levels were unaltered by the addition of talazoparib (**Fig. 6I**). Additionally, PD-L1+ and PD-1+ cells were increased with treatment, providing rationale for future testing of RT3D + PD-1 in combination with PARP inhibition (**Fig. 6J**).

## Discussion

By conducting a small-molecule drug screen, we have uncovered an unexpected role for PARP-1 in modulating RIG-I-mediated sensing and downstream signalling events in response to the presence of cytoplasmic dsRNA. We hypothesize that binding of viral dsRNA to RIG-I recruits PARP-1 which, in turn, PARylates components of the extrinsic apoptotic signalling pathway and inhibits cellular apoptosis. Initially, we postulated that this is most likely a specific property of RT3D that would allow it to maintain cell viability long enough for successful completion of its replication cycle. Instead, however, the observation that other non-viral dsRNA agonists can mediate signalling through RIG-I and recapitulate PARP-1 recruitment and inhibition of apoptosis points towards this being an important host defence mechanism to delay cell death and maintain, or even enhance, inflammatory signalling in response to a perceived viral infection. Interestingly, as anticipated, we found that pharmacological or genetic inhibition of the activity of PARP-1 led to enhanced extrinsic apoptosis pathway-driven cytotoxicity but, rather counterintuitively, an even greater degree of NFκB-mediated proinflammatory signalling. In an immunocompetent animal model, we demonstrated that the combination of RT3D and talazoparib achieved tumour control in all animals and this was associated with profound modulation of the tumour immune microenvironment and protection against subsequent isogeneic tumour rechallenge. Taken together, these findings support clinical evaluation of combinations of agonists of dsRNA sensing with PARP inhibitors and offer the prospect of developing novel treatment approaches that extend the potential utility of PARP inhibition outside the current relatively narrow confines of HR-deficient cancers.

The initial screen in A375 melanoma cells highlighted talazoparib as a strong hit, with synergistic activity at sub micromolar drug concentrations across a range of viral MOIs. This finding was confirmed in multiple melanoma cell lines representing the main genetic contexts of that disease. Contrary to our initial hypotheses, combinatorial synergistic activity was due neither to cooperative enhancement of DNA damage nor increased viral replication-related oncolysis, effectively ruling out the primary purported mechanism of action of each of the respective partners of the combination. Intriguingly, RT3D infection was shown to activate cytoplasmic PARP-1 with resulting significant PARylation of cellular proteins. As expected, PARP inhibitors significantly reduced virally-induced PARylation, and this was associated with enhanced tumour cell kill. The effect was recapitulated in PARP-1^-/-^ cells which showed exquisite sensitivity to RT3D but no combinatorial effect with PARP inhibition. Together, these data point to a role for PARP-1 in protecting cells from RT3D-induced cell death.

RT3D-induced cancer cell death involves TRAIL-mediated apoptosis and is associated with activation of Caspase-8-mediated extrinsic apoptosis^[20, 21]^. While we observed an increase in RT3D-induced TRAIL and TNF-α plus their receptors, DR4/DR5, or TNFR1/TNFR2 respectively, this was not increased further by combination with talazoparib. Immunoprecipitation for PARylated proteins pulled down DISC components (caspase-8, RIPK1 and FADD), suggesting a functional role for PARP-1 in limiting extrinsic apoptotic pathway signalling following RT3D infection alone. As further confirmation, by capturing with an anti-caspase 8 antibody, we pulled down DISC components, RIPK1 and FADD, and PARylated proteins. The addition of talazoparib markedly reduced the degree to which DISC components were pulled down by an anti-PAR antibody and this was associated, functionally, with significantly greater activity of the extrinsic apoptotic pathway. These data are consistent with previous reports of a novel function of PARP-1 in modulating caspase-8 following treatment with TRAIL, thereby inhibiting caspase-8 and limiting its function^[24]^.

Previous studies found caspase-8 can serve in two distinct roles in response to TRAIL receptor engagement: as a protease that promotes apoptosis; and as a scaffold for assembling a caspase-8-FADD-RIPK1 complex, leading to NF-κB dependent inflammation^[25]^. Furthermore, proapoptotic signals (such as Fas, TNFα, and TRAIL) were found to instigate the production of cytokines and perhaps influence immune responsiveness toward dying cells^[26]^. Similarly, our studies have confirmed that NF-κB activation occurs in response to RT3D therapy alone and that this is mediated by the DISC complex, as silencing of caspase-8, FADD or RIPK1 attenuated NF-κB and its downstream pro-inflammatory cytokines (CCL5, CXCL1, CXCL8 and CXCL10). Critically, however, we have shown that co-treatment with RT3D and PARPi markedly enhances this NF-κB-mediated inflammatory signature, despite the fact that the combination prevents the block on extrinsic apoptosis mediated by PARylation of DISC components.

The dsRNA sensor, RIG-I, has previously been shown to promote apoptosis and cell death in hepatocellular carcinoma (HCC) whilst activating NF-κB in macrophages^[27]^. Combination of RT3D plus talazoparib significantly increased RIG-I expression and massively upregulated RIG-I pathway components at the protein level, findings that correlated with the loss of PARylation. Our data point to a novel interaction between RIG-1 and PARP-1, which is amplified following RT3D plus talazoparib treatment compared to either agent alone. We posit that, following binding to dsRNA (or its mimics), the dsRNA sensor, RIG-I, can bind to and activate PARP-1 and, thus, simultaneously inhibit extrinsic apoptosis and serve as a platform for inflammatory NF-κB signalling. We further propose that combination therapy with PARPi effectively removes this PARylation-driven block on apoptosis but also functionally “traps” PARP-1 on dsRNA-bound RIG-I, in an analogous fashion to that described for PARP-trapping on DNA^[28-31]^. This leads to a phenotype of increased cancer cell death with enhanced cytokine production. Indeed, talazoparib has been shown to be a potent PARP-trapper on DNA, superior to either olaparib or veliparib. We confirmed talazoparib as a more effective and potent PARP-trapper than olaparib and veliparib (data not shown), and this correlated with greater efficacy of talazoparib in terms of triggering apoptosis, NF-κB signalling and cytokine production.

In view of the relative complexity of using replication-competent, oncolytic viral agents in the clinic, we tested if other dsRNA agonistic therapies were able to phenocopy the cytotoxic and pro-inflammatory effects of RT3D. Importantly, we confirmed that both polyI:C and 3p-hRNA were able to mediate equivalent effects and this may accelerate opportunities for early-phase clinical evaluation of this approach. PolyI:C has previously been tested in the clinic both as a poly-lysine/carboxymethylcellulose-derivatized agent^[32]^ and in nanoplexed format^[33].^ In addition, RIG-1 agonism has also been tested in a phase I clinical trial as a single agent in combination with the anti-PD1 agent, pembrolizumab^[34]^.

At this point, given the pro-inflammatory nature of the RT3D and PARPi combination, we conducted preliminary profiling in an immunocompetent BRAF^V600E^-mutant murine melanoma model. Indeed, the combination therapy was highly effective, to an even greater degree to that seen in the immunodeficient human model. All combination therapy-treated animals were cured and protected from subsequent tumour rechallenge. Importantly, RNA sequencing and flow cytometric functional analyses confirmed dramatic pro-inflammatory changes within the tumour microenvironment of animals treated with the RT3D-PARPi combination. Furthermore, these data point towards potentially effective additional combinations of dsRNA agonism, PARPi and immune checkpoint inhibition. Such approaches are the subject of ongoing work.

## Materials & Methods

### Cell lines

The following melanoma cell lines of known genetic background were used and obtained from stocks within Prof. Kevin Harrington’s team, ICR London: A375 and Mel624 (^V600E^BRAF mutant), WM266.4 (^V600D^BRAF mutant), MeWo and PWMK (wild type RAS and BRAF). D04 (N-RAS mutant) and WM17971 (K-RAS mutant) were obtained by generous donation from Prof. Richard Marais (The Paterson Institute of Cancer Research). The BRAF-mutant (BRAF^V600E^) mouse melanoma 4434 cell line was established from C57BL/6_BRAF +/LSL-BRAF^V600E^; Tyr::CreERT2+/o^[35]^ and another kind donation from Prof. Richard Marais (The Paterson Institute of Cancer Research). HeLa PARP paired cell lines (wild type PARP^++^, PARP^-/-^ [clone G3 and G9]) and HeLa clone A7 and F7 cells were obtained by generous donation from Prof. Christopher Lord, ICR London. All the cell lines were authenticated by using short tandem repeat (STR) profiling according to the manufacturer’s instructions and were carried out by our in-house sequencing facility unit. Cells were cultured in DMEM or RPMI. Media was supplemented with 5% (v/v) FBS, 1% (v/v) glutamine, and 0.5% (v/v) penicillin/streptomycin.

### Reovirus stocks

Reovirus Dearing type 3 (RT3D) stocks at 3x10^9^ tissue culture infectious dose 50 (TCID_50_/ml) were obtained from Oncolytics Biotech and stored at 1:10 concentrations in PBS at −80°C.

### Drug Screen and Cell Titer Glo (CTG) assay

Cells were plated at 500 cells per well in 20 µL, in 384 well plates using a Thermo Scientific multidrop combi and incubated overnight. An in-house drug screen (Plate 11 and 12) comprising of 80 different chemotherapeutic and targeted therapy drugs (Supplementary Table S1). We used a Hamilton microlab star robot to treat the A375 melanoma cells with doses ranging from 0.5 nM - 1000 nM. RT3D was infected into the cells (MOI ranging between 0,01-1.0) 2-hour post treatment of the drugs. Cell viability was measured by CellTiter Glo (CTG) Assay after 72 hours. After processing the CellTiter Glo luminescence data to account for plate-to-plate variation, we estimated the effect of each small molecule inhibitor on RT3D sensitivity by calculating drug effect (DE) robust Z scores, with DE Z scores <-2 being considered a profound sensitisation _effect_^[15, 16]^.

### Caspase Inhibitor experiments

The caspase inhibitor sample pack (R&D systems-Cat No FMKAP01); which contained Z-VAD (general caspase inhibitor); Z-WEHD (caspase-1 inhibitor); Z-VDVAD (caspase-2 inhibitor); Z-DEVD (caspase-3 inhibitor); Z-YVAD (caspase-4 inhibitor); Z-VEID (caspase-6 inhibitor); Z-IETD (caspase-8 inhibitor); Z-LEHD (caspase-9 inhibitor); Z-AEVD (caspase-10 inhibitor) and Z-LEED (caspase-13 inhibitor) were used at 10 μM each to inhibit the relevant caspase inhibitors in A375 and MeWo cells 8 hours prior to RT3D (MOI of 0.1) plus 0.1 μM talazoparib treatment.

### Small interfering RNA transfections

3 or 5x10^5^ cells were seeded out in the appropriate media without penicillin-streptomycin. Twenty-four hours after seeding, siRNA transfections were done on sub confluent cells incubated in unsupplemented OptiMEM using the Lipofectamine RNAi MaX reagent (Invitrogen) according to the manufacturer’s instructions. After 24 h, media was changed, and the appropriate treatment carried out for 48 hours. Lysates were collected for Western analysis, ELISA or cell death by SRB assay. All the siRNAs were purchased from Qiagen. We used two pooled siRNAs for PARP-1, 2 and 3 siRNAs: Hs PARP1_5 FlexiTube siRNA (SI02662989), Hs PARP1_6 FlexiTube siRNA (SI02662996); Hs PARP2_2 FlexiTube siRNA (SI00077917), Hs PARP2_3 FlexiTube siRNA (SI00077924); Hs PARP3_1 FlexiTube siRNA (SI00077938), Hs PARP3_4 FlexiTube siRNA (SI00077959), For the death inducing signaling complex (DISC), we used Caspase 8_11 FlexiTube siRNA (SI02661946); Hs RIPK1_5 FlexiTube siRNA (SI00288092); Hs FADD siRNA (hsFADD 7; hsFADD 5; hsFADD 8; hsFADD 9) and for the RIGI/IFN-β pathway, we used RIGI FlexiTube siRNA (GS23586).

### 3-(4,5-dimethylthiazol-2-yl)-2,5-diphenyltetrazolium bromide (MTT) assay

Cell viability was quantified using a 3-(4,5-dimethylthiazol-2-yl)2-5-diphenyltetrazolium bromide (MTT) assay. Briefly 20 μl MTT (thiazolyl blue; Sigma-Aldrich) at 5 mg/ml in PBS was added to treated cells in a 96-well plate. After 4 hours incubation at 37°C, crystals were solubilised in DMSO, and absorbance was measured at 570 nm on a SpectraMax 384 plate reader (Molecular Devices).

### Crystal violet and sulforhodamine B assays

Cell viability was quantified by staining either with crystal violet (Sigma-Aldrich) at 0.2% (w/v) in a 7% (v/v) solution of ethanol/PBS or 10% trichloroacetic acid (TCA) and stained with sulforhodamine B (SRB). The crystal violet-stained images of the plate were captured on a Microtek ScanMaker 8700 (Microtek International Ltd) while the SRB-stained cells were diluted with 1 mM TRIS and absorbance was measured at 570 nm on a SpectraMax 384 plate reader.

### Reovirus replication assays

A375, MeWo and D04 cells were seeded in 24-well plates at a density of 1 x 10^5^ cells/well. The next day cells were treated with 0.1 μM talazoparib and infected with RT3D at an MOI of 5 for 2 hours. The cells were washed twice in complete growth media. Complete growth media was added to the cells and incubated at 37°C. The cells were harvested, and the supernatants were collected at 4-, 24-, and 48-hours post-infection in triplicate. The lysates had three freeze-thaw cycles between −80°C and room temperature. For one-step growth curves the resulting lysates were titrated on L929 cells in 96-well plates. Viral titers were determined using the TCID_50_/method as previously described^[36]^. For plaque assays, dilutions were used to infect L929 cells seeded at 2 x 10^5^ cells/well in 6 well plates. After incubation at 37°C for 4 hours, the viral medium was removed and the wells overlaid with a 1:1 solution of 2% agar (Sigma) and 2X DMEM containing 5% (v/v) FCS, 1% (v/v) glutamine, and 0.5% (v/v) penicillin/streptomycin. After 5 days, plates were stained with 0.2% crystal violet in 7% ethanol. Plates containing plaques were scanned and counted using Open CFU software^[37]^. Data from the viral growth curves were derived by TCID_50_ or total viral titres quantified by plaque assays on confluent L929 cells.

### Cell cycle

Cells were seeded in 6 well dishes at 3X10^5^ cells/well and the next day treated with 0.1 μM talazoparib and/or RT3D (MOI of 0.1) accordingly. Cells were fixed with 70% ethanol at indicated time points and stained with propidium iodide (PI) at 1 μg/mL. Flow cytometry analysis was performed using a LSRII flow cytometer (BD Biosciences, Oxford, UK).

### Cell death (Celigo) Assay

Cells were seeded at 8 x10^3^ in 96 well plates and 24 hours later treated accordingly with talazoparib and/or RT3D for 48 hours post-infection. Hoechst (0.5 μg/ml) and PI (1 g/ml) were added, and the percentage of dead cells was measured using the CeligoS image cytometer (Nexcelon Bioscience).

### Confocal imaging

Cells were plated in 35 mm glass-bottomed, collagen-coated dishes (MatTek, Massachusetts, USA) and the next day treated with talazoparib and RT3D accordingly. 24 hours post-infection, cells were fixed with 4% formaldehyde and immunofluorescence performed. Cells were stained with γ-H2AX (S139) clone 20E3 (New England BioLabs, UK) or 53BP1 clone 6B3E10 (Santa-Cruz) and visualized using Alexafluor-488-conjugated goat anti-rabbit and Alexfluor-546-conjugated goat anti-mouse antibodies (Invitrogen™, Life technologies) along with 4′,6-diamidino-2-phenylindole, dihydrochloride (DAPI; Invitrogen, Molecular Probes™) nuclear stain. Cells were imaged using a LSM 710 inverse laser scanning microscope (Zeiss) and captured with a LSM T-PMT detector (Zeiss). Nuclei were quantified as positive for foci when IZ5 foci were present within the nucleus, for γ-H2AX or 53BP1.

### Western blotting

Cells were plated at 0.5 × 10^6^ in 60 mm dishes. The following day, cells were treated accordingly and collected between 24-72 hours post-treatment. Cells were washed in ice-cold PBS, pelleted and resuspended in radioimmunoprecipitation assay buffer [50 mM Tris (pH 7.5), 150 mM NaCl, 1% NP40, 0.5% sodium deoxycholate, and 0.1% SDS] with protease inhibitors (Roche Diagnostics GmbH, Mannheim, Germany), 1 mM sodium orthovanadate (Sigma), and 10 mM sodium fluoride. Cells were then lysed by snap freezing on dry ice and then allowing the lysate to thaw on ice for 10 minutes. The lysate was centrifuged at 13,200 rpm/4°C for 20 minutes to remove cell debris. Protein concentration was determined using the BCA protein assay reagent (Pierce, Rockford, IL). 30 μg of each protein sample were resolved on SDS-polyacrylamide gels (10-12%) and transferred to a polyvinylidene difluoride (PVDF) Hybond-P membrane (Amersham, Buckinghamshire, UK). Immunodetections were performed using caspase 3 [#9665], caspase 8 [#9746], RIPK1 [#3493], TRADD [#3684], DR5 [#8074], cIAP1 [#7065], RIG-I [#3743], pIRF3^(Ser^ ^396)^ [ #29047], IRF3 [#4302], pSTAT1^(Tyr^ ^701)^ [#9167], STAT1 [#9172] antibodies from New England Biolabs, UK) FADD (clone M19) [sc-6036], PARP-(clone F2) [sc-8007], PARP-2 (clone F3) [sc-393310), PARP-3 (clone B7) [sc-390771] antibodies from Santa-Cruz; PAR [4335-MC-100] antibody from Trevigen; cFLIP [AG-20B-0005] antibody from Adipogene; reovirus μ1C 10F6 and reovirus σ3 4F2 antibodies from the Developmental Studies Hybridoma Bank (IA, USA). Equal loading was assessed using α-tubulin antibody [T9026] from Sigma Aldrich. Blots were developed using secondary antibody (anti-rabbit or anti-mouse) conjugated to horseradish peroxidase (GE-Healthcare). The Super Signal chemiluminescent system (Pierce) or Immobilon Western chemiluminescent HRP substrate (Millipore) were used for detection.

### Complex II immunoprecipitation assay

Complex II purification was essentially performed as previously described^[38-40]^. Briefly cells were seeded in 10 cm dishes and treated as indicated in figure legends. After stimulation, media was removed and plates were washed with ice cold PBS to stop stimulation and frozen at −80°C. Plates were thawed, and cells lysed in DISC lysis buffer supplemented with protease inhibitors and PR619 (10 μM). Cells were lysed on ice and lysates were rotated at 4°C for 20 min and then clarified at 4°C at 13,000 rpm for 10 min. 20 μL of protein G Sepharose (Sigma) with Casp-8 (C20) antibody (Santa Cruz Biotechnology) at 1.5 μg antibody/mg protein lysate were rotated with cleared protein lysates overnight at 4°C. 4× washes in wash buffer (50 mM Tris pH 7.5, 150 mM NaCl, 0.1% Triton X-100, and 5% glycerol) were performed, and samples eluted by boiling in 50 μL 1× SDS loading dye.

### Immunoprecipitation of PARP-1

We used the PARP1-Trap Agarose kit [xtak-20] from Chromotek which consists of a PARP1 Nanobody/ VHH, coupled to agarose beads to immunoprecipitate endogenous PARP-1 proteins. Briefly, cells were harvested with Lysis Buffer: 50 mM Tris–HCl pH 7.5, 150 mM NaCl, 5% glycerol, 0.5% NP-40, 5 mM MgCl_2_, 0.2 mM CaCl_2_, 1 μM pepstatin A, 1 μM bestatin supplemented with protease inhibitor cocktail. DNase I (75-150 Kunitz U/mL) was then added to lysates and incubated for 1 hour at 4°C with rotation. Lysates were centrifuged at 4°C for 10 min at 17,000 × *g*, and cleared supernatants transferred to a pre-cooled tube. PARP-1-Trap beads were incubated with lysates for 3 hours at 4°C with rotation. Next, agarose beads were washed three times with 0.5 ml of wash buffer (50 mM Tris–HCl pH 7.5, 150 mM NaCl, 5% glycerol and 0.5% NP-40). Bound proteins were eluted with NuPAGE^TM^ LDS sample loading buffer and heated at 95°C for 5 min with 10 mM DTT.

### Proteome Profiler Human Cytokine Array

Cells were plated at 3X10^5^ cells per well in 6-well plates and media collected 48 hours after treatment with RT3D-talazoparib then centrifuged to remove cells or debris. The activity of 36 human cytokines, chemokines and acute phase proteins were simultaneously assessed using the proteome profiler human XL cytokine array kit [ARY005B] from R&D Systems (Abingdon, UK). Cell lysates were obtained from A375 following treatment with 0.1 MOI of RT3D and 0.1 μM talazoparib at 48 hours post-treatment. Cytokine secretion was carried out as a validation from the results of the human cytokine array using a range of human ELISAS from R&D Systems (see below).

### Enzyme-linked immunosorbent assay (ELISA)

Cells were plated at 2X10^5^ cells per well in 24-well plates and treated accordingly. The cells were collected at the appropriate time point and measured a range of human DuoSet ELISAs according to the manufacturer’s protocol. The following cytokines/chemokines were assessed for secretion in the cells following treatment: CCL5/RANTES [DY278], CXCL1/GROα [DY275], CXCL10/IP-10 [DY266] and CXCL8/IL-8 [DY208] and TRAIL/TNFSF10 [DY375], all from R&D systems. The TNF-α high sensitivity ELISA [BE58351] was sourced from Tecan (IBL).

### NF-κB p65 transcription factor assay

We extracted nuclear and cytoplasmic fractionation of A375 and MeWo cells following treatment according to manufacturer’s instructions (Abcam, ab113474). Briefly, cells were collected and pelleted by centrifugation then pellets were thereafter re-suspended using pre-extraction buffer from the kit. The supernatant containing cytoplasmic fraction was frozen down and stored. The pellet was centrifuged once again and re-suspended in nuclear extraction buffer from the kit to generate the nuclear fraction and samples used for the NF-κB p65 transcription ELISA based assay (Abcam, ab133112) according to the manufacturer’s instructions.

### Live cell imaging and analysis

RELA and PCNA were fluorescently tagged at the endogenous loci using CRISPR-CAS9 gene editing to generate RELA-eGFP and PCNA mScarlet-I as described on bioRxiv (https://doi.org/10.1101/2022.01.19.476961). Briefly, A375 cells were imaged at 20 min intervals following RT3D 10 MOI addition with 2 hr 1 µM talazoparib pre-treatment. Cells were imaged using the Zeiss Axio Observer Z1 Marianas Microscope with a CSUX1 confocal spinning disk unit built by 3i (Intelligent Imaging Innovations; Denver, CO). During imaging, cells were maintained at 37 °C with > 60 % humidity and 5 % CO_2._ Live imaging analysis was carried out using Nuclitrack software, which created nuclei segmentation masks from PCNA-Scarlet and quantified mean nuclear and mean perinuclear ring region levels of RELA-GFP intensity over time.

### Proteomics and LC-MS/MS Analysis

Cells were seeded in 6 cm dishes at 9 x 10^5^ cells per well overnight prior to treatments and collected at indicated time-points by trypsinising cells and washing 3 times in PBS. Pellets were stored at −80°C prior to processing. The cell pellets were lysed in 5% SDS/100 mM TEAB (tetraethylammonium bromide, Sigma) by ultrasonic probe process at 40% power for 15 x of s on / 1s off, heated at 90LJC for 10 min, and then processed by ultrasonic probe again as above. The lysate was centrifuged at 14,000 rpm for 15 min. The supernatant was collected, and protein concentration was measured by Pierce 660 nm Protein Assay (ThermoFisher Scientific) and 80 µg of proteins per sample used. Proteins were reduced by TCEP (Tris(2-carboxyethyl) phosphine, Sigma), alkylated by iodoacetamide (Sigma), and then precipitated by 20% TCA (trichloroacetic acid, Sigma) to remove detergent and excessive reagent. Protein pellet was resuspended in 80 µl of 100 mM TEAB buffer, and 3.2 µg trypsin (Pierce MS grade, ThermoFisher Scientific) was added, and the digestion went on 18 hours at 37LJC. Then 40 µg of protein digest was taken and labelled by 0.4 mg TMT11plex. 11 samples were pooled and dried in SpeedVac (Thermo scientific). The dried peptide mixture was resuspended in 0.1% NH_4_OH/100% H_2_O, and fractionated on an XBridge BEH C18 column (2.1 mm i.d. x 150 mm, Waters) with an initial 5 min loading then linear gradient from 5% ACN/0.1% NH_4_OH (pH 10) – 35% CH_3_CN /0.1%NH_4_OH in 30 min, then to 80% CH_3_CN /0.1% NH4OH in 5 min and stayed for another 5 min. The flow rate was at 200 µl/min. Fractions were collected ay every 30sec from retention time at 3 min to 44 min, and then concatenated to 51 fractions and dried in SpeedVac. The peptides were reconstituted in 30 µl of 0.1% FA/H_2_O and 50% was injected for on-line LC-MS/MS analysis on the Orbitraip Fusion Lumos hybrid mass spectrometer coupled with an Ultimate 3000 RSLCnano UPLC system (ThermoFisher Scientific). Samples were first loaded and desalted on a PepMap C18 nano trap (100 µm i.d. x 20 mm, 100 Å, 5µ) then peptides were separated on a PepMap C18 column (75 µm i.d. x 500 mm, 2 µm) over a linear gradient of 8–32% CH_3_CN/0.1% FA in 90 min, cycle time at 120 min at a flow rate at 300 nl/min. The MS acquisition used MS3 level quantification with Synchronous Precursor Selection (SPS) with the Top Speed 3s cycle time. Briefly, the Orbitrap full MS survey scan was m/z 375 – 1500 with the resolution 120,000 at m/z 200, with AGC set at 4 x 10^5^ and 50 ms maximum injection time. Multiply charged ions (z = 2 – 5) with intensity threshold at 1 x 10^4^ were fragmented in ion trap at 35% collision energy, with AGC at 1 x 10^4^ and 50 ms maximum injection time, and isolation width at 0.7 Da in quadrupole. The top 5 MS2 fragment ions were SPS selected with the isolation width at 0.7 Da, and fragmented in HCD at 65% NCE, and detected in the Orbitrap to get the report ions’ intensities at a better accuracy.

### Whole proteome data analysis

Raw spectra were processed using Proteome Discoverer v2.4(ThermoFisher Scientific) and searched against FASTA sequence databases containing GENCODE v32^[41]^ protein sequences, UniProt (2019_05) Reovirus Proteins, translated gEVE database^[42]^ sequences and cRap contaminates using both Mascot server v2.4 (Matrix Science) and SequestHT with target-decoy scoring evaluated using Percolator^[43]^. The precursor tolerance was set at 20 ppm, fragment tolerance set at 0.5 Da and spectra were matched with fully tryptic peptides with a maximum of two missed cleavages. Fixed modifications included: carbamidomethyl [C] and TMT6plex [N-Term]. Variable modifications included: TMT6plex [K], oxidation [M], and deamidation [NQ]. Peptide results were initially filtered to a 1% FDR (0.01 q-value). The reporter ion quantifier node included a TMT-11-plex quantification method with an integration window tolerance of 15 ppm and integration method based on the most confident centroid peak at MS3 level. Protein quantification was performed using unique peptides only, with protein groups considered for peptide uniqueness. Log2 fold change ratios were calculated for each sample vs time point Basal sample using normalised protein abundances. Ratios and abundance were loaded into Perseus^[44]^ or further downstream analysis and plotting. Z-score scaling was used to generate heatmaps, proteins were clustered using k-means method and GO enrichment was performed using fisher exact test. Results and RAW spectral files have been uploaded to PRIDE repository^[45]^ under project accession PXD047621.

### Quantitative RT-PCR

RNA was extracted from samples using Qiagen RNeasy kit, and cDNA synthesized using SensiFAST cDNA synthesis kit (Bioline). Samples were then amplified against transcripts by qRT-PCR with SYBR green. The following primers were used; DR4 *forward* 5-’CATCGGCTCAGGTTGTGGA-3’; *reverse* 5-’TGCCGGTCCCAGCAGACA-3’; DR5 *forward* 5-’CTGCTGTTGGTCTCAGCTGA-3’; *reverse* 5’-TGCCGGTCCCAGCAGACA-3’; TNFR1 *forward* 5-’GCTCGAGATCGAGAACGGGC-3’; *reverse* 5- ’ACGAGGGGGCGGGATTTCTC-3’; TNFR2 *forward* 5-’GGAACCTGGGTACGAGTGCCA-3’; *reverse* 5’-GCGGATCTCCACCTGGTCAGT-3’; β-actin *forward* 5’-GGCACCCAGCACAATGAA3’; *reverse* 5’-GCCGATCCACACGGAGTACT-3’. Relative gene expression was calculated by using beta actin as the housekeeping gene. All kits were used as per manufacturers’ instructions and all qpCR assays were carried out using Step One^TM^ Real-Time PCR System (Applied Biosystems).

### In vivo studies

A375 BRAF^V600E^-mutant melanoma tumours were established in female CD1 nude mice, or 4434 BRAF^V600E^-mutant melanoma tumours established in female BL6 immunocompetent mice by subcutaneous injection of 5 × 10^6^ (A375) or 4 × 10^6^ (4434) cells suspended in 100 μL PBS in the right flank. Once tumours were established and reached approximately 75-100mm^3^, mice were allocated into treatment groups stratified by tumour size before beginning therapy. talazoparib (0.1 mg/kg) or vehicle (10% DMAc, 6% Solutol, and 84% PBS) was administered by oral gavage, om day 1, 3 and 5. 1x 10^6^ pfu RT3D dissolved in PBS was administered by intra-tumoral injection after talazoparib treatment on day 3. In the CD1 models, 2 mice from each group were sacrificed on the last day of treatment then A375 tumors were harvested and homogenized in PBS on ice and extracted with Lysis Buffer [25 mM Tris (pH 7.5), 150 mM NaCl, 5 mM EDTA, 2 mM EGTA, 1% Triton X-100, sodium deoxycholate, and 0.1% SDS] with protease inhibitors (Roche Diagnostics GmbH, Mannheim, Germany), 1 mM sodium orthovanadate (Sigma), and 10 mM sodium fluoride. Caspase-8, caspase-3 and PARP cleavage were determined by Western Blot analysis while levels of PAR in the tumor lysates were determined by ELISA using the HT PARP in vivo PD Assay II Kit [4520-096-K] from R&D Systems. The remaining mice were measured twice weekly in three dimensions using Vernier calipers and the volume estimated using the formula (width x length x depth x 0.524mm^3^). In the BL6 models, 3 mice from each group were sacrificed 3 days post treatment and the 4434 tumors harvested for RNA extraction (see Gene expression analysis using Nanostring below). No toxicity or weight loss was seen in any of the treated mice. In survival studies, mice were sacrificed once tumor size reached 15LJmm in any dimension. The Kaplan-Meier survival curves were compared using the log-rank (Mantel-Cox) test using Prism Software (GraphPad). Cured mice were rechallenged on the other flank with 4x10^6^ 4434 cells and tumor growth monitored

### Gene expression analysis using Nanostring (mouse Immune Profile)

Excised 4434 tumors following treatment (as above) were lysed in homogenization tubes (Thermo Fisher Scientific) containing buffer Buffer RLT (Qiagen) using Precellys 24 homogenizer. RNA was then isolated using RNeasy Plus Mini Kit (Qiagen) according to the manufacturer’s instructions. RNA concentration was measured using Qubit Fluorometer system and Qubit RNA BR Assay Kit (ThemoFisher Scientific), and 96 ng of RNA was used in the nCounter Mouse Immunology Panel (Nanostring Technologies). Normalization, differential expression, geneset analysis and cell type scoring were performed using NanoString nSolver V.4.0 advanced analysis software. Data presentation used Rstudio V.1.4.1103, R V.4.1, ggplot2 and Complex Heatmap packages.

### Immune profiling of tumours

4434 cells were established in female BL6 immunocompetent mice by subcutaneous injection at 4 × 10^6^ cells. Tumours were allowed to grow to 6-8 mm before mice were allocated treatment groups stratified by tumour size. Treated mice were sacrificed on Day 11 and the tumors dissected and weighed. Tumors were dissociated (5 mice per group) mechanically using scissors and enzymatically digested in RPMI containing 0.5 mg/mL Collagenase type I-S (Sigma-Aldrich), 0.4 mg/mL Dispase II protease (Sigma-Aldrich), 0.2 mg/mL DNase I (Roche) and 4% Trypsin (0.25% in Tris Saline) for 30 min at 37°C. Following digestion, samples were passed through a 70 µm cell strainer and washed with 10% FCS RPMI supplemented with 5 mM EDTA. Samples were centrifuged at 1500 rpm, for 5 mins at 4°C, and transferred into a V-well 96-well plate. Samples were stained in FACS buffer (PBS + 5% FCS) for 30 mins on ice and protected from light, with the following extracellular antibodies; CD3 (100218), CD45 (103125), CD4 (100406), CD8 (562315), PD-1 (135215) from BioLegend while PD-L1 (558091) was from BD biosciences and viability dye (65-0865-14) from Thermo Fisher Scientific. Cells were then washed in FACS buffer, permeabilized and stained with intracellular antibody to FOXP3 (48-5773-80) or Ki-67 (69-5698-80) from Thermo Fisher Scientific. Samples were then washed and fixed (1-2% PFA) prior to analysis of tumour-infiltrating lymphocytes by flow cytometry. Tumours were weighed on collection and 123 count eBeads counting beads were added when running the analysis to calculate cells per mg of tumour.

### Bliss independence Analysis

Synergy interactions between different treatments were tested by standard mathematical analyses of data from MTT or SRB assays. Specifically, the presence (or absence) of synergy was quantified by Bliss Independence Analysis^[46-49]^ described by the formulae EIND = EA + EB − EA Å∼ EB and ΔE = EOBS − EIND where: EA and EB are the fractional effect of factors A and B, respectively; EIND is the expected effect of an independent combination of factors; EOBS is the observed effect of the combination. If ΔE and its 95% confidence interval (CI) are >0 synergy has been observed. If ΔE and its 95% CI are <0 antagonism has been observed. If ΔE and its 95% CI contain 0 then the combination is independent. All plots were generated using Prism GraphPad software.

### Statistical analysis

Comparisons between groups were done using the Student’s ‘t’ test or ANOVA tests. Survival curves were estimated using the Kaplan-Meier method, and significance was assessed using the log-rank test. P values <0.05 were considered to be statistically significant (*, *P* < 0.05; **, *P* < 0.01; ***, *P* < 0.005).

## Acknowledgements

We thank Oncolytics Biotech for providing reovirus and Harriet Whittock for all her help with *in vivo* work produced for this study.

