## Supplementary Figures for "PARP inhibitors enhance reovirus-mediated cell killing through the death-inducing signaling complex (DISC) with an associated NF-κB-regulated immune response"

| | 1 | 2 | 3 | 4 | 5 | 6 | 7 | 8 | 9 | 10 | 11 | 12 | 13 | 14 | 15 | 16 | 17 | 18 | 19 | 20 | 21 | 22 | 23 | 24 |
|---|---|---|---|---|---|---|---|---|---|---|---|---|---|---|---|---|---|---|---|---|---|---|---|---|
| A | | - | 5-FU | | | | sotrastaurin | | | | nilotinib | | | | AG-14699 | | | | RO-3306 | | | | + | |
| B | | - | | | | | | | | | | | | | | | | | | | | | + | |
| C | | - | carboplatin | | | | 4-OH-tamoxifen | | | | gemcitabine | | | | foretinib | | | | lapatinib | | | | + | |
| D | | - | | | | | | | | | | | | | | | | | | | | | + | |
| E | | - | BMN-673 | | | | imatinibmesylate | | | | crizotonib | | | | GSK2194069A | | | | MK2206 | | | | + | |
| F | | - | | | | | | | | | | | | | | | | | | | | | + | |
| G | | - | flavopiridol | | | | erlotinib | | | | ABT-737 | | | | methotrexate | | | | resveratrol | | | | + | |
| H | | - | | | | | | | | | | | | | | | | | | | | | + | |
| I | | - | decitabine | | | | gefitinib | | | | celecoxib | | | | MLN-4924 | | | | temozolomide | | | | + | |
| J | | - | | | | | | | | | | | | | | | | | | | | | + | |
| K | | - | compound 2 | | | | everolimus | | | | compound 1 | | | | SAR-20106 | | | | YM155 | | | | + | |
| L | | - | | | | | | | | | | | | | | | | | | | | | + | |
| M | | - | MDV-3100 | | | | compound 3 | | | | 2-methoxyestradiol | | | | cabozantinib | | | | KU0057788 | | | | + | |
| N | | - | | | | | | | | | | | | | | | | | | | | | + | |
| O | | + | sapacitabine | | | | canertinib | | | | dasatinib | | | | OSI-906 | | | | camptothecin | | | | + | |
| P | | + | | | | | | | | | | | | | | | | | | | | | + | |

| | 1 | 2 | 3 | 4 | 5 | 6 | 7 | 8 | 9 | 10 | 11 | 12 | 13 | 14 | 15 | 16 | 17 | 18 | 19 | 20 | 21 | 22 | 23 | 24 |
|---|---|---|---|---|---|---|---|---|---|---|---|---|---|---|---|---|---|---|---|---|---|---|---|---|
| A | | - | GSK-2334470A | | | | olaparib | | | | KU60019 | | | | PF-00477736 | | | | GDC-0449 | | | | + | |
| B | | - | | | | | | | | | | | | | | | | | | | | | + | |
| C | | - | BI-2536 | | | | abiraterone | | | | MK0752 | | | | PF-04929113 | | | | paclitaxel | | | | + | |
| D | | - | | | | | | | | | | | | | | | | | | | | | + | |
| E | | - | PD-0332991 | | | | lestaurtinib | | | | lenvatinib | | | | PF-332991 | | | | AZ4547 | | | | + | |
| F | | - | | | | | | | | | | | | | | | | | | | | | + | |
| G | | - | vinorelbine | | | | BIBW2992 | | | | BEZ-235 | | | | PF-00299804 | | | | XAV-939 | | | | + | |
| H | | - | | | | | | | | | | | | | | | | | | | | | + | |
| I | | - | sorafenib | | | | MK-1775 | | | | etoposide | | | | PF-03758309 | | | | GSK1904529A | | | | + | |
| J | | - | | | | | | | | | | | | | | | | | | | | | + | |
| K | | - | voronostat | | | | nutlin3 | | | | doxorubicin | | | | PF-04691502 | | | | PD173074 | | | | + | |
| L | | - | | | | | | | | | | | | | | | | | | | | | + | |
| M | | - | salinomycin | | | | BMS-911543 | | | | 6-thioguanine | | | | PF-03814735 | | | | PD-184352 | | | | + | |
| N | | - | | | | | | | | | | | | | | | | | | | | | + | |
| O | | + | sunitinib | | | | 17-AAG | | | | bleomycinsulfate | | | | PF-02341066 | | | | PLX-4720 | | | | + | |
| P | | + | | | | | | | | | | | | | | | | | | | | | + | |

**Supplementary Table 1:** Summary showing all the 80 different therapeutic cancer drugs used in the drug screen. **B**. The workflow of how the screen was done is summarised

| | *DE RT3D 0.01* | DE RT3D 0.1 | DE RT3D 0.5 | DE RT3D 1.0 | DE RT3D 5.0 |
|---|---|---|---|---|---|
| BMN-673 (1000 nM) | -1.4804556 | -4.6510624 | -7.1048246 | -10.24377 | -9.0150507 |
| BMN-673 (500 nM) | -0.101183 | -4.0332167 | -5.0411202 | -7.2730893 | -8.1487681 |
| BMN-673 (100 nM) | 0.08975608 | -2.1169224 | -3.5204909 | -6.0777899 | -4.8101895 |
| BMN-673 (50 nM) | -2.2493009 | -7.7466797 | -2.4058912 | -5.2295877 | -5.1353643 |
| BMN-673 (10 nM) | -0.6048215 | 0.75313358 | -2.7108544 | -0.3897108 | -2.0747622 |
| BMN-673 (5 nM) | -0.0707813 | 1.02504937 | -0.3610685 | -0.4173714 | -1.218146 |
| BMN-673 (1 nM) | -0.6594046 | -1.0277923 | 1.01194879 | 0.09287421 | -0.5521273 |
| BMN-673 (0.5 nM) | 0.1212299 | 0.12005849 | -0.0173739 | 0.14459598 | -1.3195996 |

**Supplementary Table 2:** Drug effect (DE) z score of RT3D and talazoparib (BMN-673)

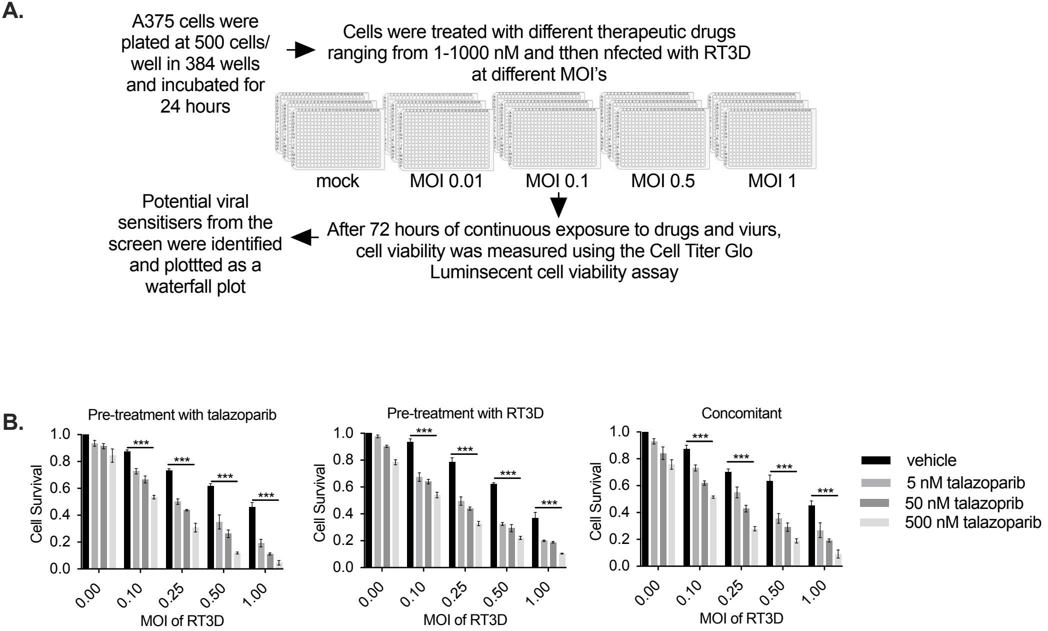


**Supplementary Figure 1: A.** Diagram summarising the setup and workflow of how the drug screen was carried out. **B**. To assess whether the dosing scheduling of RT3D *vs.* PARP inhibitor modified the sensitisation effect, A375 cells were either pre-treated with talazoparib 2 hours prior to RT3D, in a second schedule, this order was reversed and in a third schedule, talazoparib and RT3D were added concomitantly. Cell viability was assessed by MTT assay at 72 hours post–infection for all schedules. Based on these findings, we used the first treatment schedule, BMN-673 followed by RT3D, for all subsequent experiments.

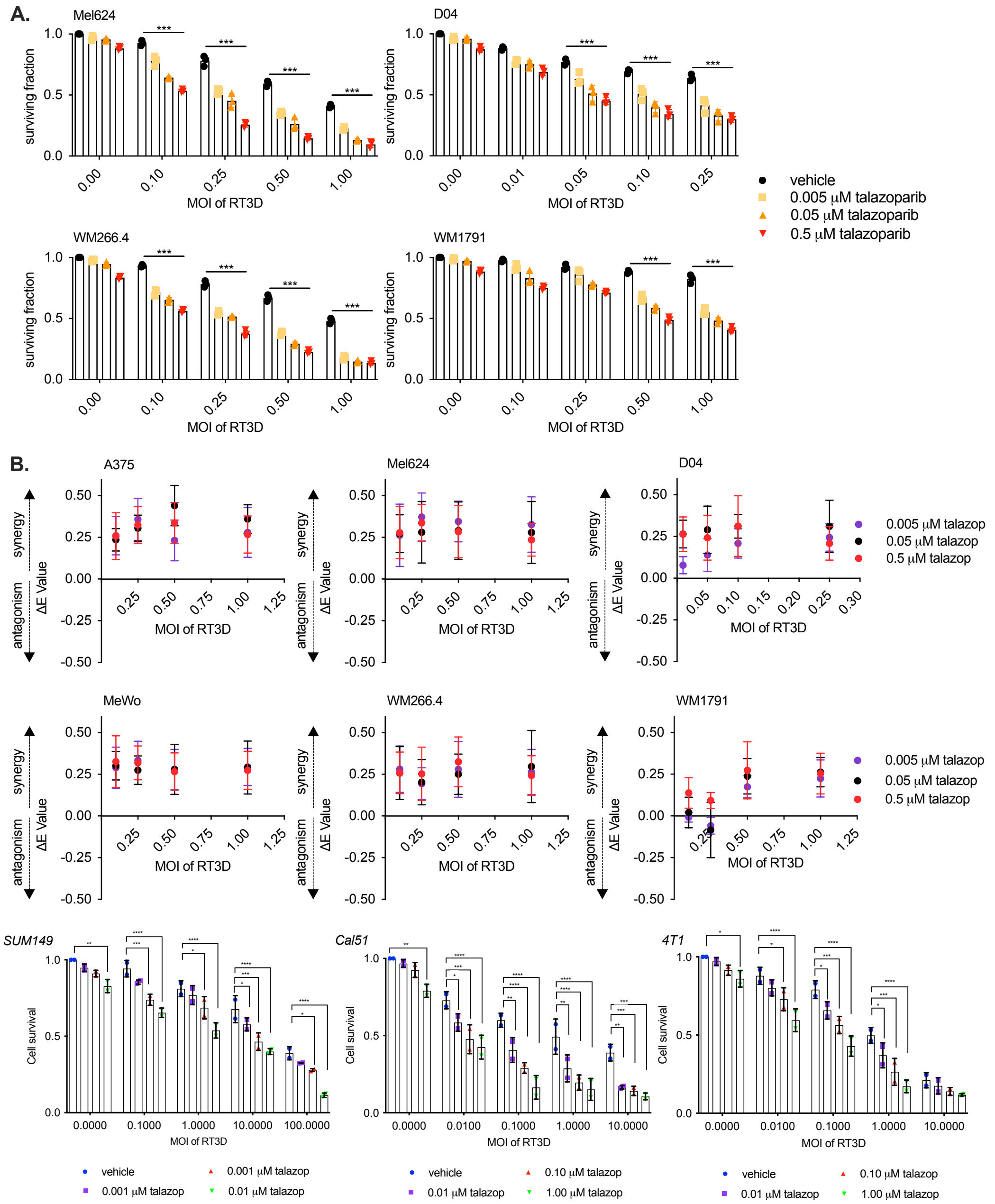


**Supplementary Figure 2: A.** A panel of melanome cells (Mel624, WM266.4, D04 and WM1791 cells) were treated with RTD and talzoparib. Cell survival was assessed by MTT at 72 hrs post-infection as shown in the bar graphs. **B.** Synergy interactions between different treatments (RT3D and talazoparib) were tested by standard mathematical analyses of data from MTT assays. Specifically, the presence (or absence) of synergy was quantified by Bliss Independence Analysis described by the formulae EIND = EA + EB – EA Å~ EB and ΔE = EOBS – EIND where: EA and EB are the fractional effect of factors A and B, respectively; EIND is the expected effect of an independent combination of factors; EOBS is the observed effect of the combination. If ΔE and its 95% confidence interval (CI) are >0, synergy has been observed. If ΔE and its 95% CI are <0, antagonism has been observed. If ΔE and its 95% CI contain 0 then the combination is independent. All plots were generated using Prism GraphPad software. **C.** A panel of triple negative breast cancer (TNBC) cell lines SUM149, Cal51 (human) and 4T1 (mouse) were treated with RTD and talzoparib. Cell survival was assessed by MTT at 72 hrs post-infection as shown in the bar graphs.

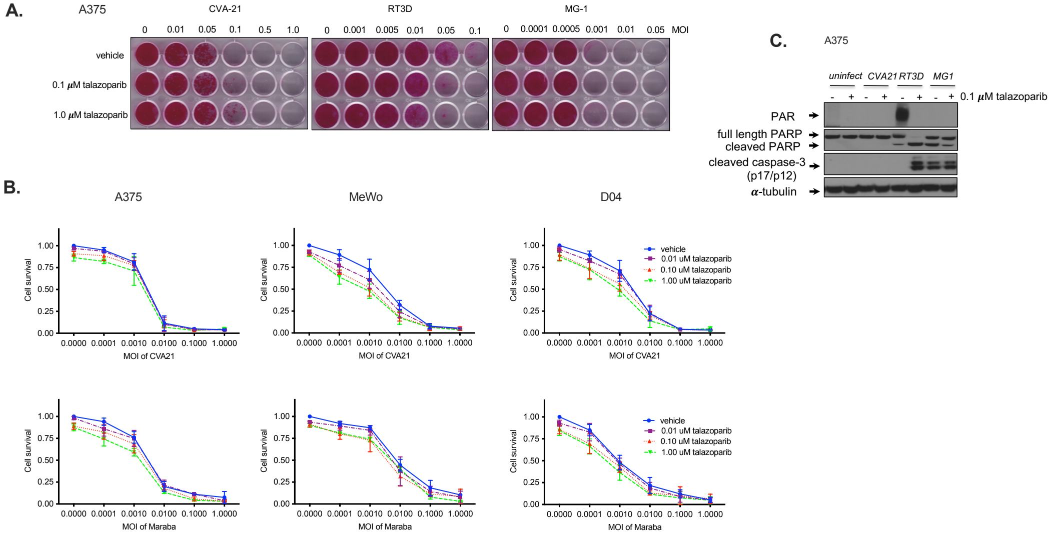


**Supplementary figure 3: A** A375 cells were treated with talazoparib at 0.1 and 1 μM in combination with CVA21 (coxsackie) MG1 (maraba) and compared with RT3D (reovirus). **B.** A panel of melanoma cells (A375, MeWo, D04 cells) were treated with talazoparib in combination with CVA21 (coxsackie), MG1 (maraba) and compared with RT3D (reovirus). Cell survival was assessed by SRB assay. C. Western analysis xxx

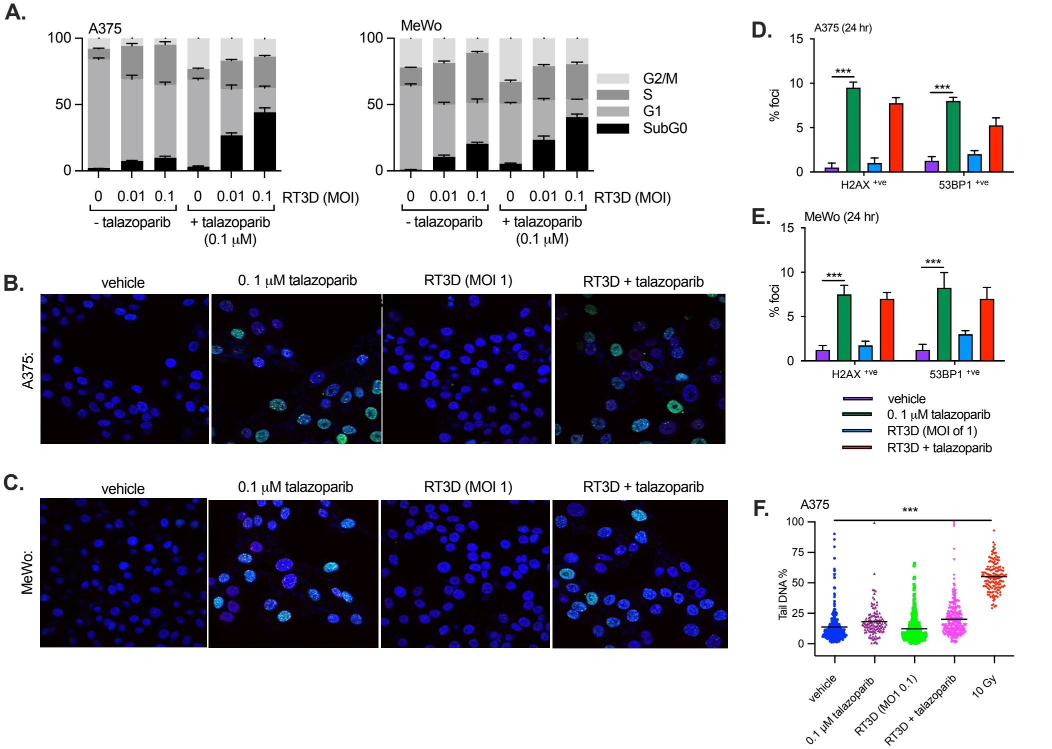


**Supplementary Figure 4: A.** Cell cycle status was assessed following with vehicle, talazoparib (0.1 μM), RT3D (MOI of 0.01, 0.1) or the combination after 48 hrs by flow cytometry using propidium iodide (PI) staining. A375 and MeWo cells were treated with vehicle, talazoparib (0.1 μM), RT3D (MOI of 1) or the combination and stained with DAPI (nucleus; blue), γH2AX (DSBs; green) and 53BP1 (DSBs; red). Images are represnted for A375 (**B**) and MeWo (**C**) at 24 hrs. The graphs measure % foci of H2AX and 53BP1 in A375 (**D**) and MeWo (**E**) cells. **F**. Comet assays were performed on vehicle, 0.1 μM talazoparib, RT3D (MOI of 0.1) or the combination treated A375 cells at 48 hrs post-treatment and 10 Gy treated sample (positive control) collected immediately after irradiation.

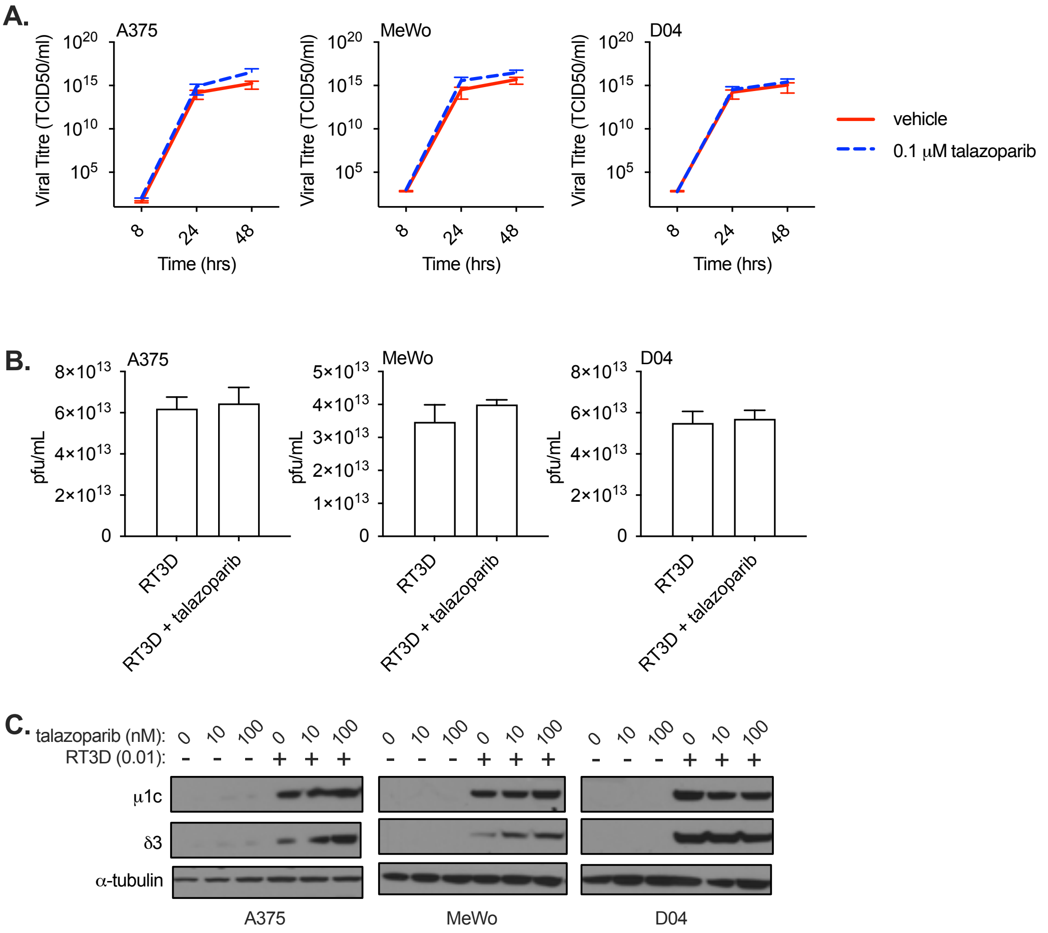


**Supplementary Figure 5:** Viral replication following RT3D and talazoparib treatment. Cells treated with RT3D plus talazoparib were harvested and the supernatants collected at 4, 24, and 48 hrs post-infection in triplicate. Viral titers were determined by using one-step growth curves (**A**), measuring the viral plaques at 48 hrs post-infection (**B**), or by analysis of viral proteins (μ1c and δ3) by western blotting at 48 hrs post -infection with RT3D at 0.01 MOI. Equal loading of proteins was assessed by probing for α-tubulin. (**C**).

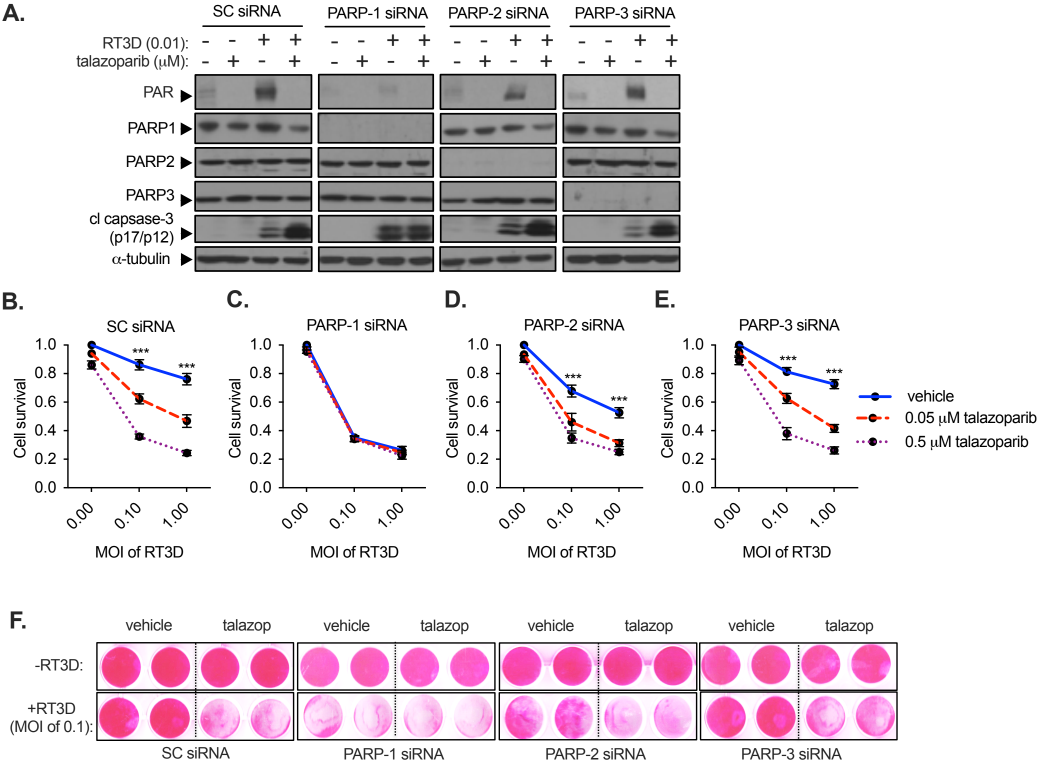


**Supplementary Figure 6**: PARP-1 plays a key role in mediating RT3D induced cell death. **A.** A375 cells were transiently transfected with scramble control (SC), PARP-1, PARP-2 or PARP-3 siRNA at 50nM prior to treatment with talazoparib (0.1μM) and infection with RT3D (MOI of 0.01). Western blotting analysis was used to probe for PAR, PARP-1, PARP-2 and PARP-3 expression. Equal loading was measured by probing for α-tubulin. Cell viability was assessed by SRB assay 72 hours following RT3D infection (0.1 and 1.0 MOI) and talazoparib (0.1μM) in A375 cells transfected with either SCR (**B**), PARP-1 (**C**), PARP-2 (**D**) or PARP-3 (**E**) siRNA. A representation of the SRB following different PARP siRNAs is shown (**F**).

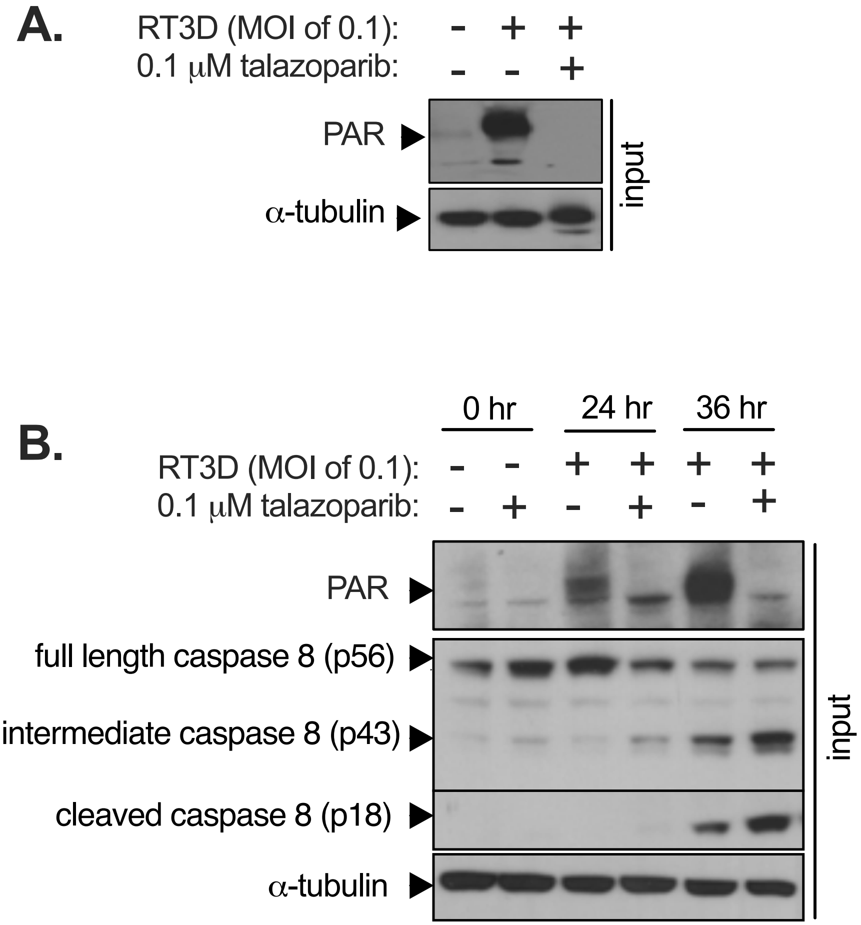


**Supplementary Figure 7**: Input (lysate) carried out to confirm RT3D induced PARylation for lysates pulled down with PAR (**A**) and pull down of caspase 8 (**B**). Equal loading of proteins was assessed by probing for α-tubulin.

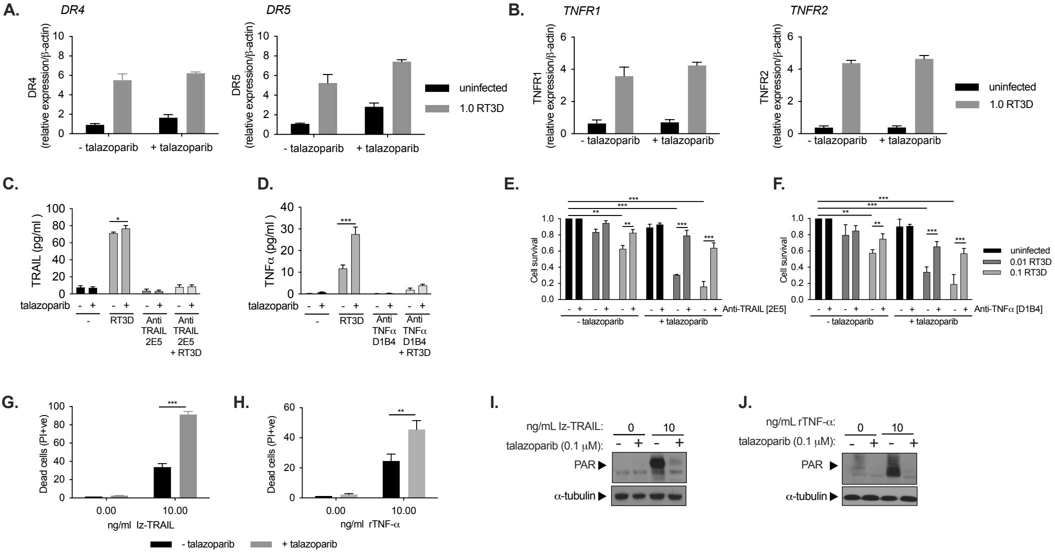


**Supplementary Figure 8**: TRAIL and TNF-α regulate RT3D/talazoparib induced cell death. **A & B**. A375 cells were pre-treated with 0.1μM talazoparib and thereafter infected with RT3D at an MOI of 1.0. DR4, DR5 (A) as well as TNFR1, TNFR2 (B) expression were validated by RT-PCR. **C & D**. A375 cells were pre-incubated with 0.2 μg/mL of Anti-TRAIL antibody (2E5) or 20 ng/mL of Anti-TNF-α antibody (D1B4) followed by treatment with 0.1μM talazoparib and RT3D (MOI of 0.1). TRAIL (**C**) or TNF-α (**D**) secretion was quantified by ELISA and confirmed loss of RT3D/talazoparib induced TRAIL or TNF-α in the presence of 2E5 or D1B4 respectively. **E & F**. A375 cells were pre-incubated with 0.2 μg/mL of Anti-TRAIL antibody (2E5) or 20 ng/mL of Anti-TNF-α antibody (D1B4) followed by treatment with 0.1μM talazoparib and RT3D (MOI of 0.01 and 0.1). Cell viability was carried out using the SRB cell viability assay following RT3D-talazoparib treatment at 48 hours post treatment and the effect of 2E5 (**E**) or D1B4 (**F**) assessed. **G & H**. A375 cells were treated with recombinant Iz-TRAIL (**G**) or TNF-α ligand (**H**) followed by talazoparib (0.1 μM). A cell death assay was used to measure the uptake of dead cells by propidium iodide (PI) 48 hours post treatment. **I & J**. PAR expression was assessed by Western Blotting analysis was carried out on A375 samples treated with recombinant Iz-TRAIL (**I**) or recombinant TNF-α (**J**) in combination with talazoparib 48 hours post treatment. Equal loading of proteins was assessed by probing for α-tubulin.

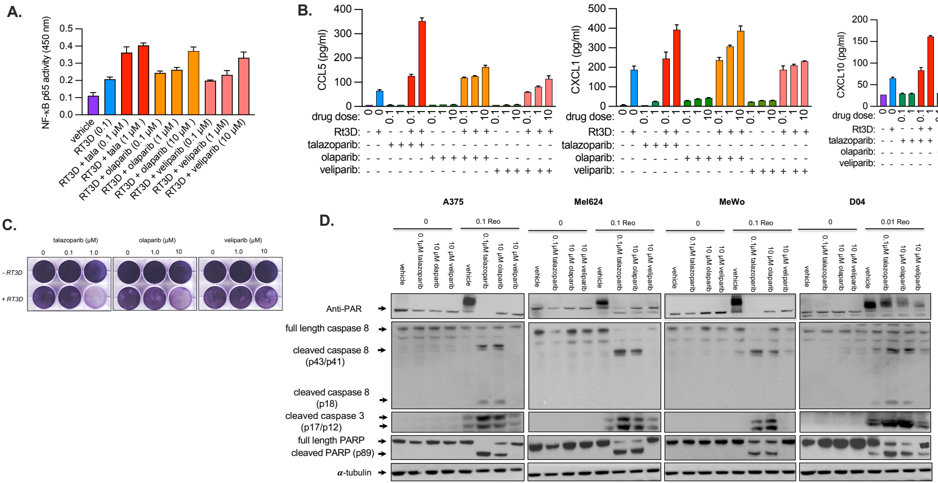


**Supplementary Figure 9**: **A.** A375 cells were treated with with either talazoparib (0.1 and 1 μM), olaparib or veliparib (1 and 10 μM) and infected with RT3D for 48 hours. NF-κB activity was measured by a colorimetric activity assay to asses DNA binding activity of the NF-κB p65 transcription factor. **B.** CCL5, CXCL1 and CXCL10 production was assessed by ELISA following RT3D infection and treatment of talazpoarib, olaparib or veliparib at various μM doses. **C.** A375 cells were treated with either talazoparib (0.1 or 1 μM), olaparib or veliparib (0.1, 1 or 10 μM) and infected with RT3D (MOI of 0.1) for 48 hours. Cell viablity assessed by crystal violet staining.. **D.** A375, Mel624, MeWo and D04 cells were treated with 0.1 μM talazoparib, 10 μM olaparib or 10 μM veliparib and thereafter infected with RT3D (MOI of 0.1) for 48 hours. Western analysis was carried out to assess PAR expression, caspase 8 and PARP cleavage. Equal loading of proteins was assessed by probing for α-tubulin.

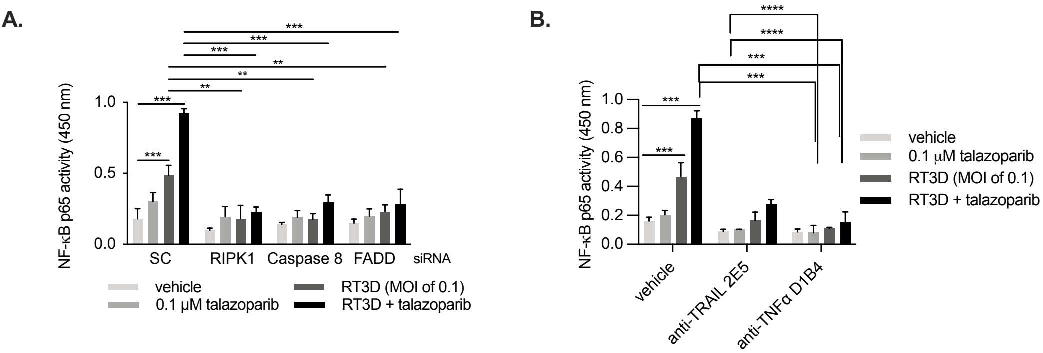


**Supplementary Figure 10**: **A.** A375 cells were treansfected with scrambled control (SC), RIPK1, Caspase 8 or FADD siRNA all at 50 nM for 24 hours and then treated with talazoparib (0.1μM) and infected with RT3D (MOI of 0.1) for 48 hours. NF-κB activity was measured by a colorimetric activity assay to asses DNA binding activity of the NF-κB p65 transcription factor RELA. **B.** A375 cells were pre-treated with Anti-TRAIL (2E5, 0.2 μg/mL) or Anti-TNFα (D1B4, 10ng/mL) for 4 hours prior to treatment with talazoparib (0.1μM) and RT3D (MOI of 0.1) for 48 hours. NF-κB activity was measured from the cell lysates by a colorimetric activity assay to asses DNA binding activity of the NF-κB p65 transcription factor RELA.

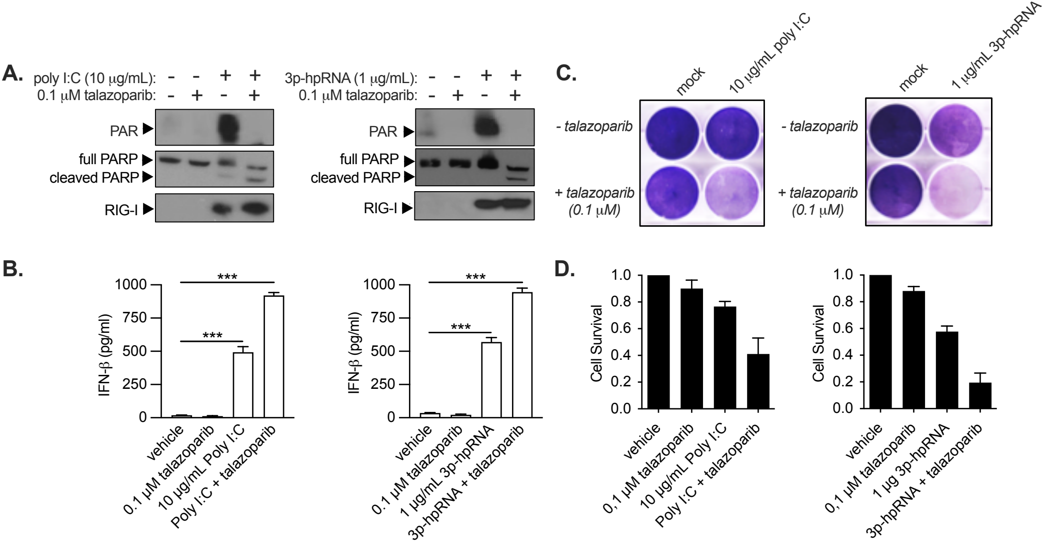


**Supplementary Figure 11**: A375 cells were transfected with 10 μg/mL poly I:C or 1μg/mL of RIG-I agonist (3p-hRNA) plus 0.1 μM talazoparib and assessed for PAR expression and PARP-1 cleavage by Western analysis (**A**), IFN-β production (**B**), crystal violet assay (**C**) and SRB assay (**D**).
